# Self-supervised predictive learning accounts for cortical layer-specificity

**DOI:** 10.1101/2024.04.24.590916

**Authors:** Kevin Kermani Nejad, Paul Anastasiades, Loreen Hertäg, Rui Ponte Costa

## Abstract

The neocortex constructs an internal representation of the world, but the underlying circuitry and computational principles remain unclear. Inspired by self-supervised learning algorithms, we introduce a computational theory wherein layer 2/3 (L2/3) learns to predict incoming sensory stimuli by comparing previous sensory inputs, relayed via layer 4, with current thalamic inputs arriving at layer 5 (L5). We demonstrate that our model accurately predicts sensory information in context-dependent temporal tasks, and that its predictions are robust to noisy and occluded sensory input. Additionally, our model generates layer-specific sparsity and latent representations, consistent with experimental observations. Next, using a sensorimotor task, we show that the model’s L2/3 and L5 prediction errors mirror mismatch responses observed in awake, behaving mice. Finally, through manipulations, we offer testable predictions to unveil the computational roles of various cortical features. In summary, our findings suggest that the multi-layered neocortex empowers the brain with self-supervised predictive learning.

## Introduction

Internal models of the external world are thought to endow the brain with the ability to predict incoming sensory information and select appropriate action-outcome contingencies ^1^. Internal models are widely believed to be encoded in the neocortex ^2,3^, whose hallmark feature is its laminar organization, comprising six distinct layers. Although much has been learned about the underlying cellular heterogeneity and connectivity of individual cortical layers, why the neocortex relies on a multi-layered structure remains unclear ^4^. Unraveling its function could shed light on the neocortical algorithms responsible for building rich internal representations of the world.

Historically, it has been proposed that *unsupervised* learning in sensory cortices underpins the development of intricate sensory representations that are critical for driving behavior ^5–7^. However, whether the laminar structure of neocortical microcircuits supports unsupervised learning remains unclear. Self-supervised learning is a form of unsupervised learning that leverages the inherent structure or patterns within the data as the target for learning. A common application of self-supervised learning is to predict the incoming input given past information ^8–12^. Importantly, self-supervised learning algorithms learn representations that better capture experimentally observed latent representations while resulting in richer models of input statistics ^12–16^. However, learning in these models is often treated as a black box, therefore it remains to be determined whether the brain is capable of employing such learning principles.

The traditional view of the neocortical microcircuit postulates a sequential flow of sensory information. In this canonical view, sensory input is relayed via the thalamus to layer 4 (L4) of the neocortex ^17,18^. L4 subsequently transmits this information to layer 2/3 (L2/3), which is thought to integrate ascending sensory information with top-down modulatory input from higher-order cortical areas ^19–21^. L2/3 in turn projects to layer 5 (L5), with L5 outputting information to other brain areas (Fig. 1a). However, growing evidence suggests that this model does not capture the full diversity of connections in the neocortical microcircuit ^18^. A body of experimental work suggests that L5 pyramidal cells receive direct thalamic input that can drive short-latency sensory evoked responses independently of activity within the cortical network (Fig. 1b) ^22–26^. These observations imply two distinct sensory-driven pathways within the neocortex, one targeting L4 and the other L5 (Fig. 1a). However, why the cortex requires multiple inputs and the computations supported by such parallel pathways remain unknown.

**Figure 1.**
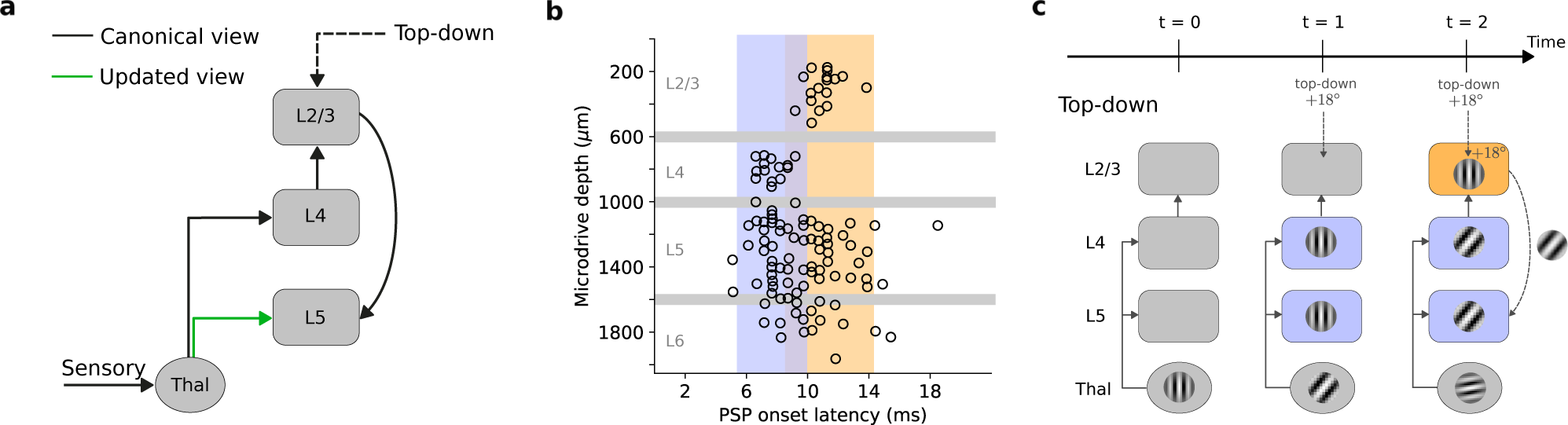
Information low in neocortical circuits. **a,** The canonical and updated view of the neocortical microcircuit. Sensory input is initially processed by the thalamus, which, in the classical view, exclusively targets layer 4 (L4). L4 subsequently relays this information to layer 2/3 (L2/3). L2/3, in turn, combines L4 input with top-down contextual input that is fed forward to layer 5 (L5). However, recent studies have emphasized the need to update this view due to direct projections from sensory thalamic nuclei to L5 pyramidal cells ^26^ (green arrow). For the sake of clarity, we omitted feedback connections from the schematic which in our self-supervised model are responsible for carrying error signals that drive learning (see main text and Methods). **b,** Onset latencies of postsynaptic potentials (PSP) by cortical depth. These results demonstrate the simultaneous activation of L4 and L5 neurons by the thalamus, indicating a direct thalamic input to L5. Adapted from Constantinople and Bruno ^26^. **c,** Proposed information flow of self-supervised temporal learning in the neocortical microcircuit. L2/3, informed by past sensory input from L4 and top-down contextual input, predicts the current sensory input arriving in L5. The direct thalamic inputs to L5 provide sensory input, which is used as a teaching signal to instruct the L2/3 predictive model.

Inspired by this refreshed view of the canonical microcircuit and the predictive capabilities of self-supervised machine learning algorithms ^9,27^, we propose a model in which L2/3, informed by past sensory input from L4 and top-down context, predicts incoming sensory input. In this model, the L4-to-L2/3 delay enables L2/3 to generate predictions based on previous sensory information. Direct thalamic input to L5 is responsible for providing sensory-based teaching signals required for L2/3 to adjust and refine its predictions (Fig. 1c). This perspective of the neocortical circuitry suggests that the L4-L2/3-L5 laminar structure with parallel thalamic innervation enables the brain to learn rich temporal representations.

We first show that our learning rule for L2/3-to-L5 connections closely resembles experimentally observed long-term synaptic plasticity ^28^. Using self-supervised learning, our model can learn and predict Gabor-like inputs in contextual sequential tasks, highlighting its effectiveness in capturing complex patterns. By ablating individual components of the model and evaluating their impact on performance, we reveal how the neocortical circuit components collaboratively enable self-supervised learning. Next, we demonstrate that self-supervised learning leads to predictions that are robust to sensory noise and occlusions. Moreover, the model predicts layer-specific sparsity that closely matches those found experimentally in sensory systems. Additionally, we demonstrate that our model produces L2/3 and L5 prediction errors in response to visuomotor mismatches, thereby providing an explanation to the mismatch responses observed in awake, behaving animals. Finally, we suggest a set of optogenetic experiments capable of testing the core predictions of our self-supervised learning model. Collectively, our findings support the notion that the L4*→*L2/3*→*L5 pathway is instrumental in enabling the brain to engage in temporal self-supervised learning, underscoring its potential significance in neural mechanisms of predictive learning.

## Results

### Neocortical layers can implement self-supervised predictive learning

To understand how neocortical microcircuits process temporal information and learn latent representations in a self-supervised manner, we created a model that emulates the properties of cortical circuits. Our model contains three subnetworks with non-linear neurons separated into L2/3, L4, and L5 to reflect the laminar architecture of the neocortex (Fig. 2a). Within this framework, L4 receives ascending sensory information, *x*, at timestep *t* through input weights *W*_Thal._*_→L_*_4_. L2/3 receives delayed input, timestep *t−*1, from L4 via *W_L_*_4_*_→L_*_2_*_/_*_3_ synapses and top-down contextual input via *W*_top-down_*_→L_*_2_*_/_*_3_ weights. We hypothesize that this combination of inputs enables L2/3 to make predictions about upcoming sensory information. L2/3 predictions are then sent down to L5 via *W_L_*_2_*_/_*_3_*_→L_*_5_ synapses and compared with the actual sensory input received by L5 at timestep *t*. This comparison enables an error function to be computed which is then fed back via L5-to-L2/3 connections to adjust the L2/3 predictive model of incoming input. The error function is defined as 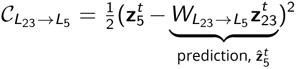 where 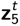 is the activity of L5 neurons and 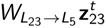 its prediction, 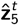. In addition to this predictive error, L5 is also trained to reconstruct its own input (see Methods). During learning, we modify connections to minimize the error functions and facilitate the encoding of sensory input. Consequently, our model requires both feed-forward connections from L4 *→* L2/3, which relay sensory information, as well as feedback connections from L5 back to L2/3, to transmit error signals. All weights are optimized via gradient descent.

**Figure 2.**
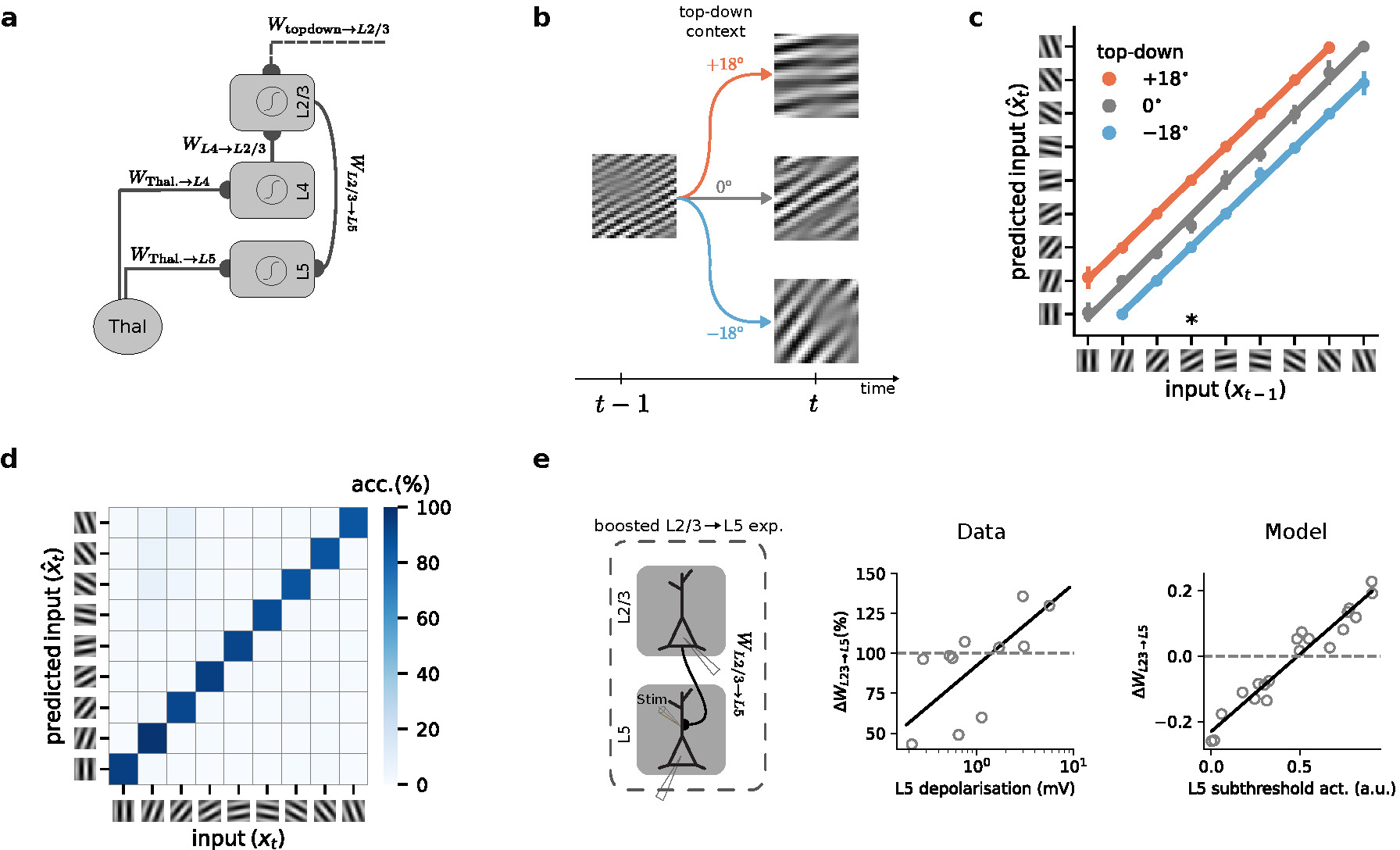
A model of temporal self-supervised learning in cortical circuits. **a,** Schematic of the cortical circuit model. **b,** Schematic of a sequential Gabor task. The generative factor that is provided to the model as top-down context at timestep *t* determines the orientation of the next Gabor patch at timestep *t* + 1. **c,** Decoding accuracy of a linear model trained on the output of L2/3. For a given input, L2/3 predicts the incoming sensory input with high accuracy. Colors represent the three possible conditions (*−*18*^◦^*, 0*^◦^*, and +18*^◦^*). *∗* points to the example illustrated in panel b. **d,** Confusion matrix for classification accuracy of a linear model trained on the output of L5. The metrics in c and d are calculated over 5 different initial conditions. **e,** Left: schematic of the experimental setup in which an extracellular electrode was used to boost L5 activity while inducing long-term synaptic plasticity on L2/3-to-L5 connections ^28^. Middle: observed changes in synaptic weights as a function of L5 depolarization (scatter plot: individual data points, solid line: linear fit to the data). Right: L2/3-to-L5 learning rule as predicted by our model as a function of L5 activity for multiple randomly drawn samples of L2/3 and L5 activity (circles), and linear fit to the data points (solid line). Error bars represent the standard error of the mean over 5 different initial conditions.

To demonstrate our model’s ability to learn useful representations, we created two-step sequences of Gabor patches, which are commonly used to evoke responses in the primary visual cortex (see Methods) ^29,30^. Starting with a random Gabor patch, at time point *t*, the orientation of the Gabor patch changes according to top-down contextual input that is randomly generated (see Methods). This higher-order contextual cue to L2/3 is provided at each time step and mimics signals such as those provided by the motor cortex ^31^. Given this top-down input the Gabor patch either rotates anti-clockwise by *−*18*^◦^*, remains the same, or rotates clockwise by +18*^◦^* (Fig. 2b). For example, if the current Gabor patch has 0*^◦^* orientation, the orientation of the subsequent input can be *−*18*^◦^*, 0*^◦^*or +18*^◦^* for contextual cue values of *−*18*^◦^*, 0*^◦^*, and +18*^◦^*, respectively (Fig. 2c).

To evaluate the representations learned by our model, we use linear decoders ^8,32,33^. We trained this decoder on L2/3’s output and compared it with the Gabor patches received by L5 at a given timestep. L2/3 effectively learns to predict upcoming Gabor patches using the previous input, provided by L4, and the top-down context value (Fig. 2c). Next, we applied a linear classifier on L5’s output. On average, L5 achieves a test accuracy of *≈* 89% (classification accuracy on a random model is *≈* 11%), indicating that L5 successfully identifies and encodes each Gabor patch’s distinct features (Fig. 2d). In addition, we obtain similar results in a more complex task in which hand-written digits are used as input (Fig. S1). Our results indicate that L2/3 achieves near-perfect accuracy (*≈* 93%) in predicting the subsequent Gabor patch, outperforming L5. These results are in line with experimental findings showing that L2/3 can learn to predict image sequences ^30^.

Next, we compared the learning rule for the key predictive weights in our model, *W_L_*_2_*_/_*_3_*_→L_*_5_ (Methods Eq. 7), with observed long-term synaptic plasticity in primary sensory cortices (Fig. 2e) ^28^. Our learning rule predicts a depression-to-potentiation switch as the activity of L5 neurons increases. This is in line with experimental observations showing a similar depression-to-potentiation switch of *W_L_*_2_*_/_*_3_*_→L_*_5_ connections with increasing depolarization of L5 pyramidal cells ^28^. Hence, this experimental evidence corroborates our model’s learning rule, showing that the model is consistent with the updated view of the neocortical circuit and known synaptic plasticity mechanisms in primary sensory cortices.

In summary, we have shown the model’s ability to perform self-supervised learning in a temporal task and its consistency with synaptic plasticity observations. However, we have yet to explore the precise contribution of each circuit element to self-supervised learning.

### Neocortical circuitry jointly underlies self-supervised learning

In our model, different cortical layers give rise to distinct computational roles. To demonstrate this, while generating experimentally testable predictions, we systematically ablated individual connections, allowing us to quantify their impact on both representational capability and performance (Fig. 3A).

**Figure 3.**
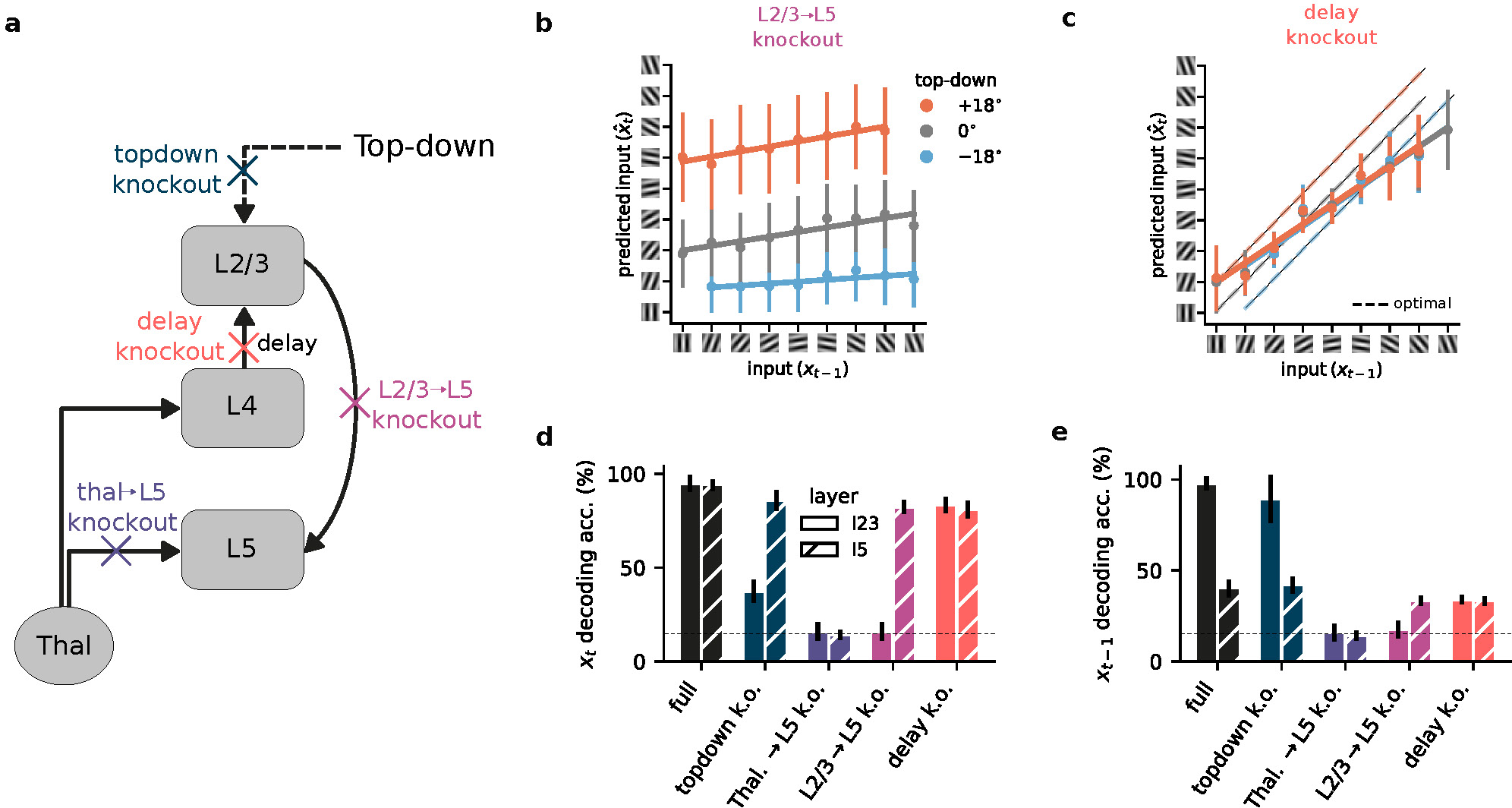
Neocortical circuitry jointly enables self-supervised learning. **a,** Schematic of the model with individual components knocked out (colored crosses) within the neocortical microcircuit architecture. **b,** Connections from L2/3 to L5 are necessary for L2/3 to learn a predictive representation of the input. **c,** Impact of L4-mediated delay in self-supervised learning (dashed lines represent the optimal prediction). **d,** Summary of decoding accuracy of the current input for L2/3 and L5 when specific connections are knocked out. The x-axis indicates the specific ablation, while the y-axis the decoding accuracy for the current input (*x_t_*). **e,** Similar to d, but for the past input (*x_t−_*_1_). Knockout components are color-coded as in panel a. Horizontal dashed lines in d and e represent chance decoding accuracy. Error bars represent the standard error of the mean over 5 different initial conditions.

When we knocked out the L2/3-to-L5 connection, L2/3 was unable to learn to make predictions about upcoming sensory information (Fig. 3b vs. Fig. 2c). This is due to the lack of communication between the source of the prediction in L2/3, and the source of error signals in L5. As a consequence, the L2/3 predictive model does not learn. However, because top-down contextual input is still present, L2/3 still shows contextual segregation.

Our model proposes a key function for the delay introduced by L4 as information propagates to L2/3. This delay creates a temporal discrepancy between the information available to L2/3 (past input) and L5 (present input) which enables L2/3 to learn predictive representations by attempting to anticipate the incoming sensory input. When this delay is removed, which would be equivalent to L2/3 receiving direct thalamic input, the entire network operates on incoming sensory input, i.e. at timestep *t*. Consequently, the network can no longer generate meaningful predictions of future inputs. This is reflected in Fig. 3c where L2/3 fails to reliably distinguish among potential future outcomes. This result highlights the key role that temporal delays may have in shaping predictive learning within the neocortical microcircuit. The L4 to L2/3 delay is essential for biasing L2/3 representations towards the future. Without it, both the Thal. *→* L4 *→* L2/3 *→* L5 and Thal. *→* L5 pathways end up representing the current sensory input, rendering the former pathway redundant.

Next, we investigated how the ablation of these different circuit elements affects the ability to decode *current* sensory information from L2/3 and L5 representations. For current input decoding (Fig. 3d), L5 demonstrated robust accuracy as long as it retained access to thalamic sensory input. This aligns with its role as a primary recipient of sensory data, together with L4^23,26,34^. L2/3 accuracy, however, was more dependent on the circuit properties. While top-down input to L2/3 provided useful context-dependent input (Fig. S2), any disruption to core pathways within the microcircuit, except the delay knockout, compromised L2/3’s ability to represent the current sensory input.

Decoding the *previous* input (Fig. 3e) further differentiated L2/3 and L5. As anticipated, L5 exhibited limited information about previous inputs due to its exclusive focus on current thalamic information. L2/3, however, encodes information about the past as a result of the delay introduced by L4. Complete loss of this past-input representation occurred only when critical learning pathways were ablated (Thal. *→* L5, L2/3 *→* L5), or when the delay was removed; thereby synchronizing L2/3 and L5 inputs.

Our ablation and decoding analyses suggest that predictive learning within the neocortical microcircuit depends on a complex interplay between L2/3 and L5. We find that the L2/3-to-L5 connection is essential for the model to learn predictive representations, suggesting that L2/3 prediction errors drive learning. Furthermore, the temporal delay between L4 and L2/3 is crucial for generating future-oriented predictions, but not current representations. In terms of decoding past and present sensory input, our results demonstrate that L2/3 specializes in representing temporal context, while L5 primarily encodes immediate sensory information. This result aligns with the experimental observations showing that L2/3 effectively encodes temporal information with high precision ^35^.

### Self-supervised learning leads to robustness to noise and occlusion in cortical networks

As a consequence of learning a robust predictive model of the sensory input, the network should disregard unpredictable aspects like noise. Therefore, we hypothesized that the self-supervised L2/3 *→* L5 predictive component would help L5 filter out sensory noise. To test this, we ablated the L2/3 *→* L5 projection in our model. This simulated ablation shows that removing the predictive component (i.e. *L*2*/*3 *→ L*5) dramatically reduces robustness to different noise levels (Fig. 4a; similar results are obtained when ablating the L4-to-L2/3 delay, Fig. S3). This denoising capability emerges naturally from the model’s design, even though it was not explicitly designed for this purpose, which are comparable to a near-optimal denoising autoencoder network.

**Figure 4.**
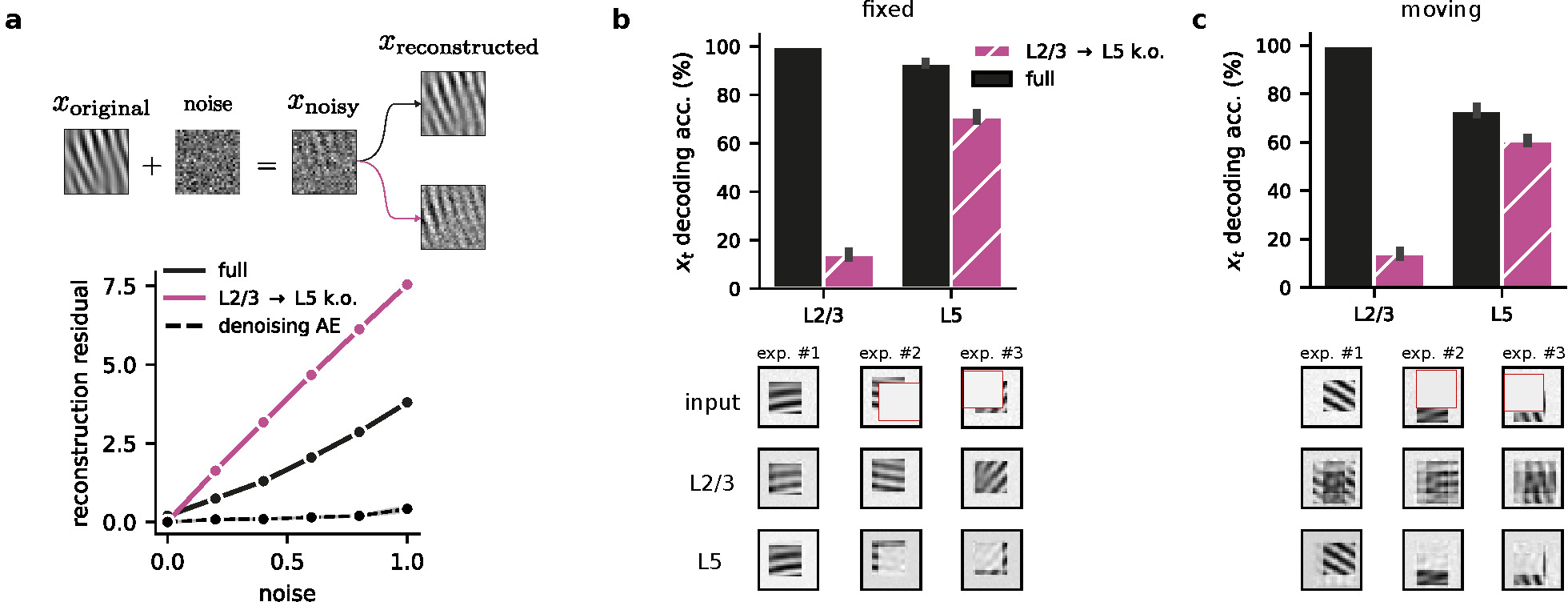
L2/3-to-L5 predictions are crucial for denoising and resolving occluded stimuli. **a,** L2/3-to-L5 connections promote noise suppression in L5 representations. Top: Schematic of noise added to the original inputs. Bottom: Noise-corrupted input samples lead to higher L5 reconstruction residuals (*x̂_t_ − x_t_*) when the L2/3*→*L5 pathway is ablated (purple) compared to the full model (solid black). We also provide the reconstruction residual for an autoencoder that was explicitly trained to denoise the input (dashed black line). **b,** Top: Decoding accuracy with and without L2/3*→*L5 for a Gabor task with occlusion. Bottom: Three examples depicting L2/3’s ability to recover occluded information, compared to L5’s incomplete reconstructions (top row: original occluded input; middle row: L2/3 prediction; bottom row: L5 reconstruction). **c,** Top: Accuracy with and without L2/3*→*L5 connections for a task in which Gabor patches move randomly. Bottom: Examples illustrating the robustness with moving Gabor patches (top row: original input with motion (cf. panel b); middle row: L2/3 prediction; bottom row: L5 reconstruction). L2/3 reconstruction encodes uncertainty about the future possible input location. Error bars represent the standard error of the mean over 5 different initial conditions.

To further investigate the model’s robustness to input perturbations we tested its ability to reconstitute input patterns during partial input occlusions. We modified our sequential Gabor task, by randomly occluding parts of the input (Fig. 4b). After training the model without occlusions, we assessed the robustness of the learned representations in each layer by evaluating their ability to classify occluded sensory input. We observed that L2/3 achieves higher decoding accuracy compared to L5 and that L5 decoding is not affected by L2/3 *→* L5 knockout (Fig. 4b, top). Further analysis shows that L2/3 can fully reconstruct the input while L5 is only able to reconstruct the observable parts of the input (Fig. 4b, bottom). These results support the idea that a strong predictive model leads to representations that are robust to several perturbations.

Finally, we explored whether L2/3 can also encode the uncertainty about the possible input locations. To test this, we introduced random shifts in the position of the Gabor patch on the blank canvas during the task. Decoding performance remained similar to the fixed-position task, but reconstructions were different (Fig. 4c, top). L2/3 representations reflect the input’s positional uncertainty (blurred reconstructions across possible locations), while L5 again encodes only the visible parts (Fig. 4c, bottom). This suggests that self-supervised learning also leads to useful L2/3 representations when in the presence of sensory uncertainty.

Collectively, these results underscore the robustness exhibited by the proposed neocortical predictive learning model across diverse input conditions. Consequently, our model offers valuable insights into the mechanisms through which cortical circuits deal with the considerable variability inherent in naturalistic environments.

### Layer-specific sparseness emerges from self-supervised learning

Sparse coding, in which only a small subset of neurons are strongly active for a given stimulus, is a widespread phenomenon across the neocortex ^36–40^. This sparsity is particularly pronounced in superficial layers (L2/3) compared to deeper layers (e.g. L5) ^41^. However, it is unclear why the degree of sparsity varies across cortical layers and how it relates to their computational role.

We wondered whether our network, equipped for temporal self-supervised learning, could reproduce experimentally observed sparsity distributions. Moreover, we wanted to investigate how different network features may control sparsity across different neocortical layers. To this end, we trained our model on the sequential Gabor task. After training, we measured population sparseness across layers using established metrics ^42^ (see Methods). Interestingly, our results closely mirror experimental findings ^41^: L2/3 presents the highest sparseness, followed by L4 and then L5 (Fig. 5a). This alignment suggests that self-supervised learning, focused on input prediction, could be a key factor driving sparsity in biological neural networks.

**Figure 5.**
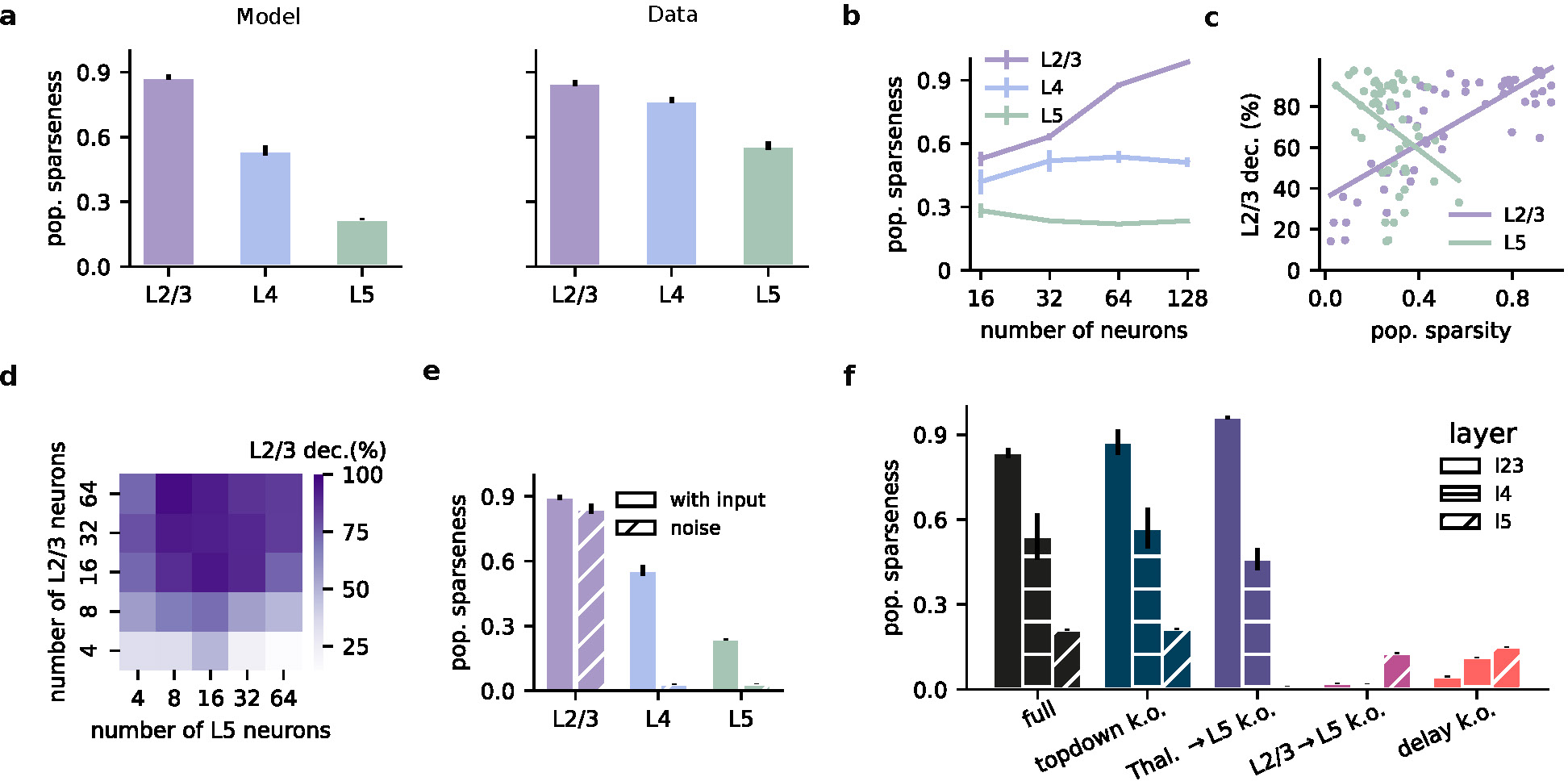
Population sparseness depends on neocortical layer. **a,** Population sparseness across layers in the model (left) and experimental data ^41^ (right). **b,** Population sparseness as a function of the number of neurons. The qualitative relationship between layers is preserved, but L2/3 sparseness increases with network size. **c,** Decoding accuracy of current input as a function of the population sparsity of L2/3 and L5 (L2/3: r=0.78, p=2.1e-11; L5: r=-0.35 p=0.01). **d,** L2/3 decoding accuracy of the current input as a function of the number of neurons in L2/3 and L5. **e,** Population sparseness with or without sensory input (noise condition) after learning. L2/3 remains sparse, while L4 and L5 show a strong reduction in response sparsity. **f,** Population sparseness following ablation during learning of different model components. Top-down input ablation slightly increases L2/3 sparseness. Thalamic input ablation to L5 decreases L2/3 sparseness while increasing L5 sparseness. Ablation of L2/3-to-L5 connections abolishes sparseness across all layers. Error bars represent the standard error of the mean over 5 different initial conditions.

Layer 2/3 has undergone rapid expansion relative to other layers within the human evolutionary lineage ^43,44^. Could this expansion support greater predictive learning capabilities? We found that L2/3 sparseness increased with network size, while sparsity in L4 and L5 remained relatively stable (Fig. 5b). Consistent with the increased sparsity in L2/3, we find that an increase in the number of L2/3 neurons also results in improved L2/3 decodability of upcoming sensory inputs (Fig. 5c,d), in line with previous work ^45^. In contrast, the relationship between the number of neurons, sparsity, and decoding accuracy was not present in L5 neurons (Fig. 5c,d; Fig. S4).

To determine whether sparsity is due to the encoding of sensory input, or is simply an underlying feature of the circuitry, we replaced sensory input with Gaussian background noise. When sensory input was replaced with background noise, L2/3 retained high sparsity, whereas L4 and L5 responses showed a strong decrease in response sparsity (Fig. 5e). Importantly, this is an effect that emerges over learning (Fig. S4). These results suggest that L2/3 sparsity is primarily a consequence of learning to predict sensory input, whereas deeper layers rely more heavily on ongoing input dynamics.

Next, we ablated different model elements during learning to test their contribution to the emergence of response sparsity (Fig. 5f). Removing top-down input had a minimal effect on sparseness. However, ablating thalamic input to L5 during learning selectively decreases L5 sparseness, which is likely due to the resulting random L5 responses. Finally, ablating L2/3-to-L5 connections or the delay component completely abolished sparsity across all layers, demonstrating their crucial role in encouraging sparseness over learning.

Together, these results show that sparsity emerges as a function of input-driven predictive learning as postulated by our model, thus providing an explanation for layer-specific sparsity as observed experimentally.

### L5 *→* L2/3 feedback is required for self-supervised learning

A cardinal feature of self-supervised learning models is that they require an error, or teaching signal to instruct plasticity across the network. This error signal prompts adjustments in the synaptic weights, thereby refining the network’s activity to enhance its predictive model. Since the error signal that drives learning in our model originates in L5, the resulting error signals should, in principle, be transmitted back to L2/3 pyramidal neurons to improve the L2/3 predictive model. Therefore, this suggests the need for a feedback connection that propagates this information from L5 to L2/3. Although the vast majority of work on neocortical circuits has disregarded feedback connections from L5 to L2/3 pyramidal cells ^17,18^, growing evidence shows that they are more abundant than previously assumed ^46,47^.

Here we explore the importance of the L5 to L2/3 feedback connection for learning in the model. In particular, we contrast *optimal* feedback, as used in previous figures, with *random* and *no feedback*. The optimal feedback condition corresponds to a setting in which the feedback weights mirror the feedforward weights (i.e. 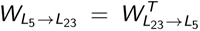), whereas in the random feedback condition the feedback weights are set to a random weight matrix ^48^.

Inspired by work showing that random feedback weights are suffcient for credit assignment ^48^, we tested whether this form of unstructured feedback was suffcient for L2/3 to learn. We observed that L2/3 was indeed able to learn to predict the input with random feedback weights (Fig. 6a), in line with the optimal feedback (cf. Fig. 2c). These results suggest that unstructured feedback may be suffcient in enabling L2/3 to develop useful predictive representations. Furthermore, our findings demonstrate a significant drop in the decoding accuracy of L5 when feedback connections from L5-to-L2/3 (Fig. 6b) were removed. This decline is due to L2/3’s inability to learn, causing L5 to adopt erroneous representations influenced by L2/3’s unlearned state. These results demonstrate the need for L5 *→* L2/3 feedback.

**Figure 6.**
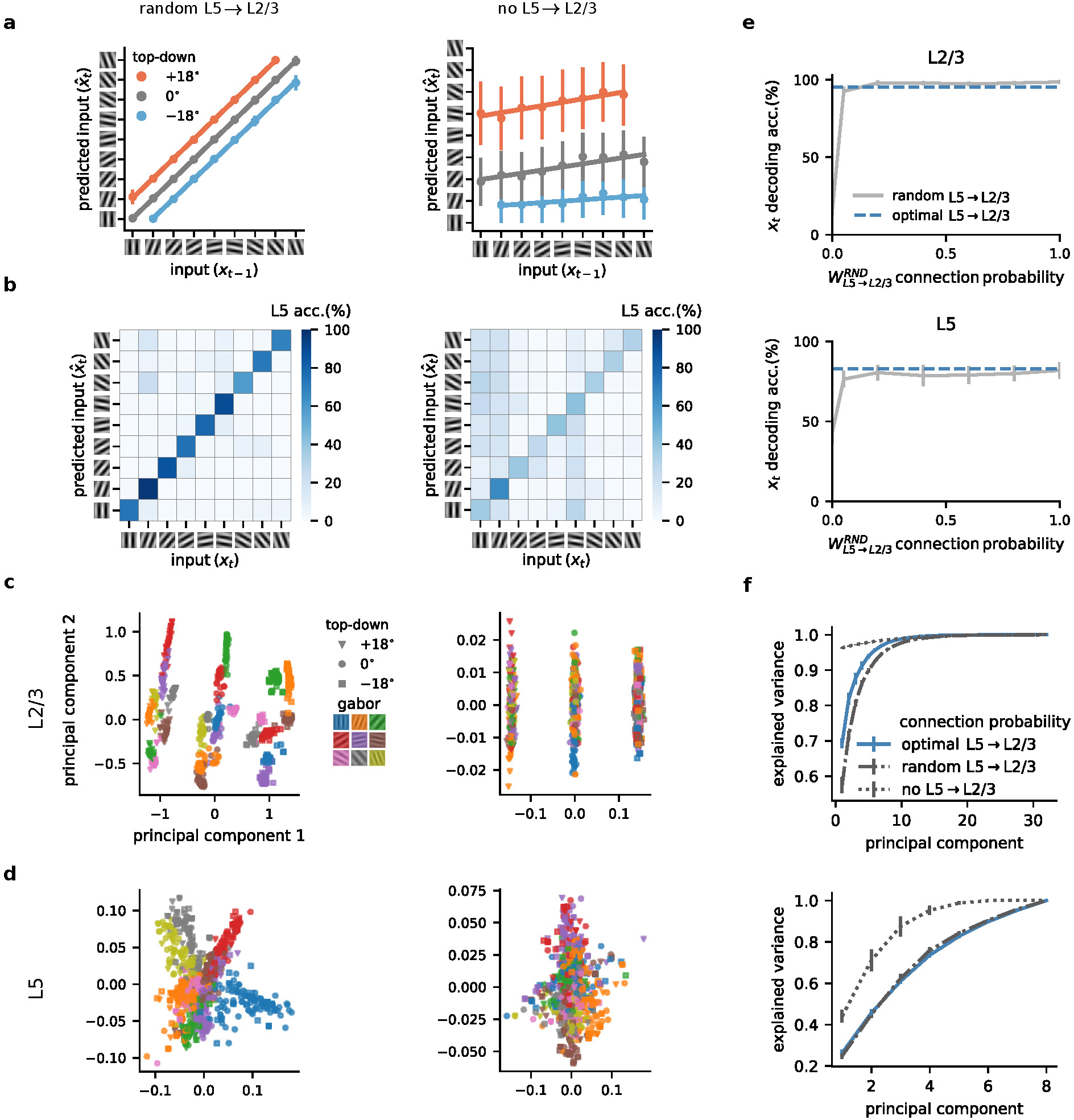
Role of L5-to-L2/3 feedback connections in self-supervised predictive learning. **a,** L2/3 learns to predict the input in the presence of random feedback (left) but fails to do so without L5-to-L2/3 feedback (right). **b,** L5 learns to represent the inputs accurately with random feedback (left) but shows lower decodability without feedback (right). **c,** Two main principal components of layer 2/3 representations for random (left) and no feedback (right) across different top-down contexts (symbols) and input Gabor orientations (colors). **d,** Two main principal components of layer 5 representations for random (left) and no feedback (right) across different top-down contexts (symbols) and input Gabor orientations (colors). **e,** L2/3 (top) and L5 (bottom) decodability for different degrees of L5-to-L23 feedback. **f,** Explained variance of L2/3 (top) and L5 (bottom) learnt representations. Error bars represent the standard error of the mean over 5 different initial conditions.

Next, to study the neuronal representations learned by the different layers, we analyzed the two main principal components of L2/3. This revealed a notable difference in the structural organization across feedback conditions. With feedback, L2/3 representations were differentiated based on the identity of the Gabor patch as well as the top-down context (Fig. 6c, left). Without feedback, the L2/3 representations were only distinguished based on top-down inputs, indicating a limitation in the network’s learning capability (Fig. 6C, right).

Similar analysis of L5 showed that with random feedback, representations are grouped as a function of Gabor patches, suggesting a structured learning process (Fig. 6d, left; similar to the optimal feedback condition, Fig. S5). These observations are in line with the increased sparsity we observed in L2/3 compared to L5 (Fig. 5). In contrast, when feedback was absent, L5 representations were less organized (Fig. 6d, right).

Classically, feedback connections within the neocortex occur at lower probabilities than the corresponding feed-forward pathway ^46,47^. To test how connection density influenced the properties of the network, we next explored how the linear decoding accuracy of both L5 and L2/3 varies with the probability of feedback connections from L5 to L2/3. An increase in connection probability corresponded to enhanced decoding accuracy (Fig. 6e). However, while a very low feedback connection probability was suffcient for learning the task considered here, for more complex tasks higher connection probabilities may be required for optimal performance (Fig. S6).

Finally, to determine how distributed information was, we examined how the explained variance, as assessed by the number of principal components (PCs), changed with varying feedback probabilities. The absence of feedback required a greater number of PCs to explain the data effectively, while random feedback closely mirrored the effciency of optimal feedback connections (Fig. 6f). This increase in the number of PCs to capture the same variance is consistent with our findings above, showing the importance of feedback in organizing sensory information in superficial layers (cf. Fig. 6c,d).

This analysis underscores the critical role of feedback connections in neural networks, particularly in enhancing predictive capabilities and structuring neural representations. The nuanced differences observed across varying feedback types and intensities offer insights into the role of observed L5-to-L2/3 feedback connections ^46,47^ in learning and information processing in neural networks.

### Model generates sensorimotor prediction error signals consistent with experimental observations

Our study thus far evidences the capability of our cortical model to make predictions about upcoming sequences. We next sought to determine how the model responds when those predictions are violated and if these responses differ for superficial and deep cortical layers. We also wanted to test whether our network generates prediction error signals that resemble those observed in cortical networks of behaving animals ^19,31,49^. For example, recent *in vivo* awake experiments have observed error-like mismatch responses in a visuomotor task ^19,49^.

We aimed to test if our model could reproduce the mismatch responses in both L2/3 and L5 recently observed experimentally ^19^. In the study by Jordan and Keller ^19^ the authors explored the prediction error responses in a setting where animals learn to couple the speed of the visual flow (that is, sliding vertical gratings) to speed of locomotion (Fig. 7a). This paradigm allows for the systematic investigation of how neural responses in the primary visual cortex are shaped by the interplay of external sensory stimuli (visual flow) and internal contextual expectations (running speed). Using whole-cell recordings, Jordan and Keller showed that when the visuomotor coupling is temporarily broken (i.e. ‘visuomotor mismatch’), the majority of L2/3 pyramidal neurons depolarize, whereas the majority of L5 excitatory neurons hyperpolarize. We propose that the opposing mismatch responses between L2/3 and L5 observed experimentally ^19^ can be explained by visuomotor prediction errors in a network implementing self-supervised predictive learning.

**Figure 7.**
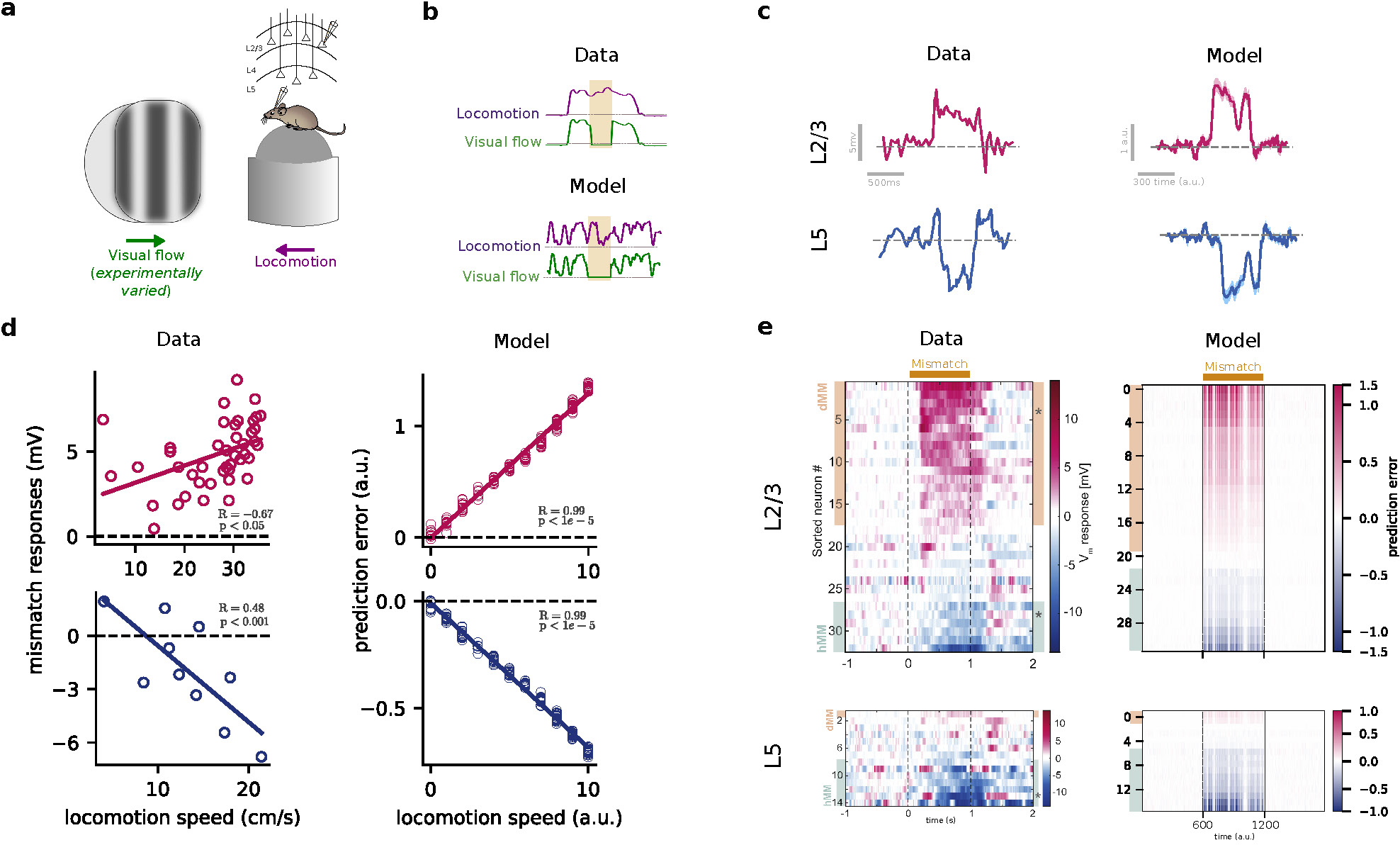
Model generates sensorimotor mismatch prediction errors in line with experimental observations. **a,** Illustration of visuomotor task used by Jordan and Keller ^19^ in which mice learned to associate visual flow with locomotion. **b,** Sample of training data from experiments (top) and our synthetic dataset (bottom). As in the experimental setup, we randomly halt the visual flow (flat green line) to generate visuomotor mismatches. **c,** When the visual flow is halted, a sample neuron in L2/3 of the mouse visual cortex shows depolarisation, while a sample neuron in L5 shows hyperpolarization (left). In our model, in line with the data, L2/3 shows a positive mismatch error, while L5 shows a negative mismatch error (right). Shaded areas represent standard deviation over 5 runs. **d,** The mismatch error signals in the model are correlated with the modelled locomotion speed when the visual flow is halted (right), in line with experimental observations ^19,49^ (left). **e,** Model generates a distribution of mismatch prediction errors which are biased towards positive errors in L2/3 and negative errors in L5 (right), in line with mismatch responses observed in primary visual cortex ^19^ (left). Schematics in (a) and data in (b), (d), and (e) were adapted from Jordan and Keller ^19^, Padamsey and Rochefort ^50^.

To this end, we produced a synthetic dataset that captures the correlation between visual flow and running speed. Specifically, running speed is dictated by a random walk process whereby the speed at any given instance is determined based on the preceding speed plus a random variation (see Methods). Under normal conditions, changes in visual flow were linked to changes in speed by linearly scaling the visual flow vector according to the running speed. Occasionally, we uncoupled visual flow and locomotion by setting the visual flow to zero, thus generating visuomotor mismatches (Fig. 7b). With this setup, we can investigate how the network responds to both intact and uncoupled visuomotor integration.

We first explored the sign and magnitude of prediction errors in our model. To this end, we calculated L2/3 and L5 prediction errors using the gradients of the self-supervised error function with respect to L2/3 and L5 neurons, respectively (see Methods). When visual flow was randomly halted during locomotion, we observed a positive error signal in L2/3 neuronal activity and a negative error signal in L5 (Fig. 7c). This discrepancy arises from their roles, as postulated by our self-supervised learning model: L2/3 predicts the upcoming visual flow while L5 encodes the actual visual flow. Hence, when visual flow halts and given top-down running speed, L2/3 predicts a non-zero flow, thereby resulting in a positive prediction error. L5, in contrast, encodes the actual zero flow, and, hence, generates a negative prediction error due to the (positive) prediction provided by L2/3 input. Both L2/3 and L5 mismatch errors in our model are consistent with mismatch responses observed experimentally (Fig. 7c). Furthermore, we found that the mismatch responses in our model scale linearly with the running speed (Fig. 7d), in line with experiments ^19,49^.

To further test whether the mismatch errors across neurons resemble those that have been found experimentally ^19^, we analyzed the distribution of prediction errors predicted by our model. Our results show that the majority of L2/3 neurons exhibit positive mismatch responses while the majority of L5 neurons exhibit negative mismatch responses (Fig. 7e; see also Fig. S7 for prediction errors across different model conditions), again in line with Jordan and Keller ^19^.

### Differential role of L2/3 and L5 during sensorimotor mismatches

We have shown that the prediction-error responses observed in L2/3 and L5 of awake behaving animals ^19^ can be explained by the different roles played by cortical layers equipped with self-supervised predictive learning. This suggests that layer-specific manipulations, for example through simulated optogenetic stimulation, of L2/3 and L5 neurons may reveal their unique functions. To test this in our model, we selectively increase the neuronal response of either L2/3 or L5 neurons during sensorimotor mismatches to determine their contributions to the prediction errors in the other layer, L5 and L2/3, respectively.

To this end, we first scale the output of L5 neurons. Upon moderate scaling, the the model predicts that both positive and negative mismatch errors in L2/3 should decrease in magnitude (Fig. 8b). By increasing the L5 activity further, the neurons that displayed a positive mismatch error before scaling, now switch their sign and become negative mismatch error neurons. Similarly, the neurons that displayed a negative mismatch error before scaling, now signal a positive mismatch error (Fig. 8a,b). These results are a consequence of L5 neurons representing zero input during the sensorimotor mismatch period while L2/3 neurons predict a non-zero visual flow as a result of nonzero topdown. By increasing the L5 output activity, eventually, the feedback signal from L5 becomes larger than the prediction in L2/3, which leads to a change in the signs of prediction errors in L2/3.

**Figure 8.**
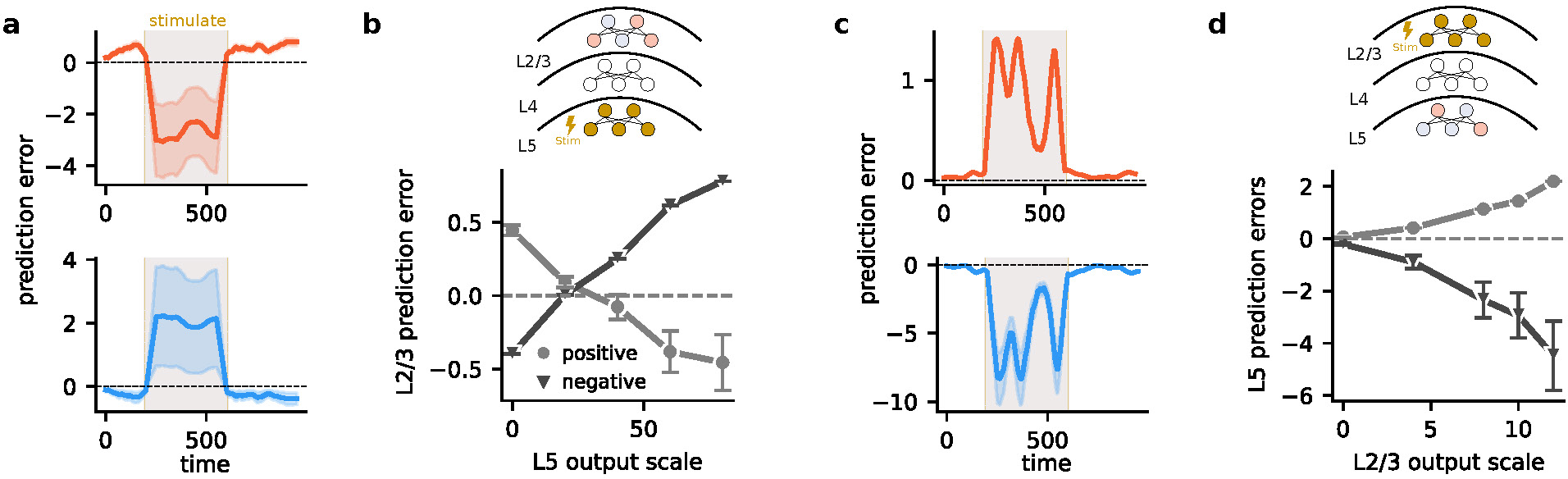
Differential role of L5 and L2/3 activation in generating mismatch prediction errors. **a,** Stimulating L5 during the mismatch interval causes prediction errors in L2/3 to switch signs. Neurons with positive errors switch their error signal with scaling of L5 activity (top) and vice versa (bottom). **b,** Positive errors gradually shift towards negative and vice versa, demonstrating a direct relationship between L5 stimulation and L2/3 prediction error modulation. **c,** Stimulating L2/3 during the mismatch interval amplifies existing prediction errors within L5. **d,** Plot demonstrates a proportional increase or decrease in prediction error magnitude as the output of L2/3 is scaled.

Thus far, we have scaled the L5 output altogether, without distinguishing between neurons exhibiting a positive or negative mismatch error. When we scaled only the outputs of neurons exhibiting a positive mismatch in L5, the changes in L2/3 mismatch errors were heterogeneous (Fig. S8). This is likely due to the low number of neurons with positive mismatch errors in L5, limiting their impact on L2/3. In contrast, scaling only the output of the L5 neurons exhibiting negative mismatch errors reverses the mismatch errors in L2/3 (Fig. S8), similar to the impact of scaling the L5 output altogether (Fig. 8b).

Next, we studied how changes in L2/3 affect the mismatch errors in L5. Scaling the output of L2/3 neurons altogether enhances the mismatch errors in L5. That is, neurons exhibiting a negative mismatch error before scaling now show even stronger negative mismatch errors after L2/3 activity scaling. Likewise, neurons exhibiting a positive mismatch error before scaling, now exhibit even stronger positive mismatch errors after scaling (Fig. 8c,d). This remains true even when we only scale the neurons with a positive mismatch error in L2/3 (Fig. S8). However, scaling only the L2/3 neurons with a negative mismatch error reverses the mismatch errors in L5 (Fig. S8). Hence, our model predicts a dynamic interplay between L2/3 and L5, with L2/3 stimulation generally amplifying L5 mismatch errors. Interestingly, this effect is primarily driven by L2/3 neurons signaling positive mismatch errors, while stimulating the negative mismatch neurons in L2/3 reverses the direction of prediction errors in L5.

These results can be best explained by the asymmetric contribution of L5 and L2/3 in generating mismatch errors. The strong effect of neurons with a positive mismatch error in L2/3 aligns with their role in predicting sensory input; increasing their activity strengthens this prediction signal within L5. Conversely, neurons with a negative mismatch error in L2/3 likely represent neurons suppressed by a greater-than-expected input. Enhancing their activity during a mismatch further emphasizes this “less than expected” signal, ultimately reversing the sign of the prediction error in L5.

Overall, these targeted manipulations of layer-specific mismatch signals offer a means by which to dissect the distinct functional roles of L5 and L2/3 populations. This approach could be further explored experimentally to advance our understanding of how neocortical layers contribute to predictive learning of sensory streams.

## Discussion

Inspired by a refreshed view of the canonical neocortical circuitry and modern self-supervised learning algorithms, we introduce a computational theory wherein L2/3 learns to anticipate incoming sensory input. We demonstrated L2/3’s capacity to predict incoming sensory information using temporal-contextual tasks. As a result, L2/3 develops latent sensory representations that are resilient to sensory noise and occlusions, improving the ability of cortical networks to encode partially observable information. Additionally, the proposed optimization leads to layer-specific sparsity, in line with experimental findings. Subsequently, by employing a sensorimotor task, we reveal that the model’s prediction errors align with L2/3 and L5 mismatch responses observed in awake, behaving mice. Finally, using manipulations we generated predictions for the role of specific circuit elements in self-supervised predictive learning.

Our study focuses on the canonical L4-L2/3-L5 three-layered motif ^17,18^. This classical view of the neocortical microcircuit emphasizes the feedforward flow of information across layers. However feedback projections are also evident ^46,47^ and both anatomical and electrophysiological data suggest the existence of direct thalamic input onto L5 pyramidal cells that effectively bypasses this feedforward circuity ^22,23,25,26,51^. Our model explores the computational significance of these pathways by mapping them onto a self-supervised learning framework. Our results suggests that the two parallel thalamic pathways play critical, yet distinct roles: the L4-L2/3 pathway makes a temporal prediction, while the thalamic-L5 pathway provides the self-supervised target (i.e. incoming sensory input) with which to test this prediction. Feedback from L5-to-L2/3 connects the two parallel systems to guide learning.

Our model proposes that a critical feature of L2/3’s integrative capacity is to use past information to predict incoming sensory input. This aligns with experimental observations showing that superficial layers display stronger temporal integration of sensory information compared to deep layers ^52,53^. Our work indicates that the delay introduced by the thalamic-L4-L2/3 pathway is critical for the emergence of these properties. It also predicts that the delay introduced by these neurons and synapses sets the predictive time scale achievable by L2/3 neurons. Since this delay is approximately 10-20 milliseconds ^26,54^, this suggests that the temporal horizon for prediction in L2/3 of primary sensory cortices is limited. However, the cortex is highly hierarchical. Our framework, combined with higher-order top-down inputs, may provide a circuit-based explanation for why higher-order brain areas show longer timescales ^55,56^. For example, the secondary visual cortex (V2) also gets primary thalamic projections, indicating that V2 could introduce further delays in the sensory input via cortico-cortical projections onto the L4*→*L2/3 pathway. These delays could then be compared with the incoming sensory input in L5, allowing the brain to learn predictive representations over hundreds of milliseconds or even longer.

It has been well documented that superficial layers respond more sparsely to sensory stimuli than deep layers ^41^. However, it was not known how this feature emerges. In our model, layer-specific sparsity occurs naturally due to the proposed predictive function of L2/3 (Fig. 5). Our results also imply that simple measures like sparsity can help infer the optimization/learning processes of various brain structures. Indeed, we show how selective ablations of individual components of the network alter sparsity in a layer- and pathway-specific manner. While these findings provide important insights into the nature of sparsity in cortical networks, fully understanding the interplay between input, connectivity, and the emergence of sparsity during learning requires further investigation.

While our model has been mapped onto the canonical six-layered structure of the neocortex, it operates with only three layers. This raises an intriguing consideration: the evolutionarily conserved three-layered structures found in other brain regions, such as the hippocampus and piriform cortex in mammals, as well as in the cortices of other species like turtles ^57^, may represent the foundational blueprint for self-supervised learning. This suggests that a three-layer structure might serve as the fundamental building block for self-supervised learning, which was subsequently elaborated upon throughout evolution, firstly by increasing the number of layers and then secondly by expanding the primary locus of self-supervised learning in L2/3. Consistent with this view, we also find that sparsity increases as the size of L2/3 increases (Fig. 5b). L2/3 is greatly expanded in human evolution, even when compared to other layers ^43,44^. Our results indicate that such expansion may improve network function, including expanding the predictive capabilities of the human neocortex.

In our model, we derive prediction errors directly from optimization principles. Consequentially, our model predicts the need for feedback between neocortical layers that carries information about error signals. This provides a form of credit assignment within the neocortical microcircuit. Although models of the neocortex often disregarded feedback pathways, numerous experimental studies demonstrate their existence ^46,47,58^. Despite this, these pathways are understudied and their organization is not well understood, in contrast to feedforward which often shows highly organized subnetwork architectures ^59–61^. Our findings indicate that both structured, i.e. reciprocal, as well as sparse, random feedback enable learning, with the former potentially advantageous for more complex tasks. A further question is how feedback error signals may be computed in a biologically plausible manner. Recent work shows how this can be achieved using dendritic compartments and interneuron cell types ^62–64^. Different interneuron sub-types, with distinct connectivity, control feedforward and feedback processing in the L2/3-L5 circuitry ^65–68^, which may underlie distinct aspects of the self-supervised learning proposed here.

While these studies suggest biological mechanisms through which self-supervised learning may emerge, is there evidence that it occurs within the brain? Recent experimental studies support the ability of the neocortex to perform self-supervised learning ^69^, while deep networks trained using self-supervised learning better capture experimentally observed representations compared to networks trained via supervised learning ^13^. For example, training deep networks using self-supervised predictive error functions yields representations that resemble visual cortical features ^8,16,70–73^. Taking a step towards understanding the underlying learning mechanisms, recent research has introduced a combination of Hebbian and predictive synaptic plasticity ^12^. This body of work supports the notion that sensory cortices engage in self-supervised learning, yet the specific circuit-level computations facilitating this process have remained unclear. Our work ties self-supervised learning to specific neocortical layers, suggesting that L2/3 and L5 provide complementary roles for implementing self-supervised learning. Consistent with these findings, the L2/3-to-L5 pathway is highly conserved across cortical regions ^74,75^ and behavioral studies have highlighted its importance for learning ^76^. In future work, it would be of interest to test our theory by performing layer-specific experiments ^77,78^.

Predictive coding has provided a framework by which to learn sensory representations in the brain ^79–81^. It postulates that the brain learns an internal model of the world from sensory streams by directly updating neuronal dynamics through prediction errors ^1,7^. In temporal predictive coding, the neural networks constantly attempt to predict the incoming stimulus. Lotter et al. ^82^ demonstrated that a deep convolutional network that is trained using predictive coding learned sensory representations useful for downstream tasks. This contrasts with our model, where prediction errors lead to plasticity, rather than changes in neuronal dynamics directly. Temporal predictive coding models are also often relatively abstract and do not consider how predictive coding is implemented. A notable exception is the work of Bastos et al. ^1^ in which it was proposed that L5 encodes input expectation while L2/3 encodes positive and negative prediction errors in separate populations. This is in contrast with our model in which L2/3 predicts the incoming input while L5 encodes the current sensory input and computes prediction errors locally, which in turn explains a range of experimental observations. A feature of our model which goes beyond existing predictive coding models is the fact that our model jointly predicts the incoming sensory input in L2/3 together with representing the current input in L5, in line with recent developments in deep learning ^83^. In future work, it would be of interest to explicitly contrast our model with existing predictive coding frameworks.

Overall, our work suggests that circuit motifs found throughout the neocortex implement effcient self-supervised predictive learning in the brain.

## Materials and Methods

We model the dynamics of the neocortical circuitry using a set of connected neuronal layers. The architecture consists of distinct layers corresponding to the L4, L2/3, and L5 layers of the neocortex. The connectivity between layers is all-to-all unless otherwise stated.

In our model, L4 neurons receive direct thalamic input, *x_t_*, and their activity is given by

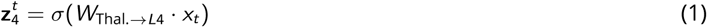

where *σ* is the sigmoid function and *W*_Thal._*_→L_*_4_ is the weight matrix that models the connectivity from the thalamus to all L4 neurons.

We then model the neuronal and synaptic delay introduced by L4 as information flows onto L2/3 by one timestep. This means that L2/3 integrates past inputs from L4 (i.e. from time step *t −* 1) with top-down inputs from higher-order cortical areas at time step 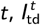. L2/3 is modelled as,

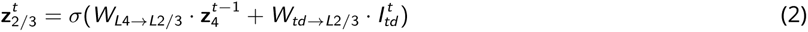

where **z**_23_ is a vector with all neurons in L2/3 and *W_L_*_4_*_→L_*_2_*_/_*_3_ is the weight matrix from L4 to L2/3. As above, all neurons are subject to the sigmoid non-linearity *σ*. L5 receives direct thalamic input and L2/3 input. It is modeled as,

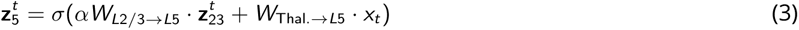

where **z**_5_ is a vector with all L5 neurons, *W_L_*_2_*_/_*_3_*_→L_*_5_ and *W*_Thal._*_→L_*_5_ are the weight matrices from L2/3-to-L5 and thalamus-to-L5, respectively. *α* is a constant that models the dendritic-to-somatic attenuation of L2/3-to-L5 input. We set *α* = 0.3, but the exact value does not qualitatively change our results.

In our network, the weight matrices *W_L_*_2_*_/_*_3_*_→L_*_5_, *W_L_*_4_*_→L_*_2_*_/_*_3_, *W*_Thal._*_→L_*_4_, *W*_Thal._*_→L_*_5_ and *W_td→L_*_2_*_/_*_3_ are subject to optimization through gradient descent. The learning rules for these connections follow cost/error functions inspired by those commonly used in machine self-supervised learning. In particular, we use a combination of two cost functions,

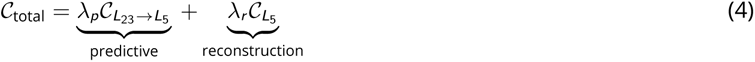

*λ_p_* and *λ_r_* are hyperparameters that scale the predictive and reconstruction costs, respectively. The first component of *C*_total_ is the temporal self-supervised cost in which L2/3 *predictions* given L4 input at time *t −* 1 are compared with L5 inputs at *t*

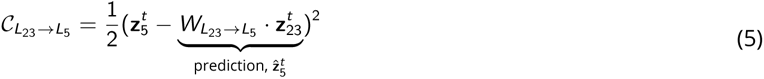

The second component of *C*_total_ encourages the model to learn non-trivial representations through *reconstruction* of L5 thalamic input given its own activity, as follows

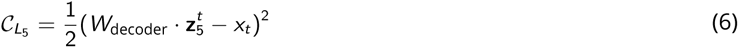

This reconstruction cost is only used to optimise *W*_Thal._*_→L_*_5_ and *W*_decoder_ weights. To ensure this, we block the resulting error signals (i.e. gradients) from adjusting *W_L_*_2_*_/_*_3_*_→L_*_5_ weights and the remaining weights.

The key predictive learning rule is the one that governs 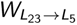 weights. It can be derived from the total cost as,

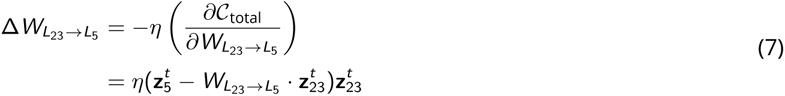

where *η* denotes the learning rate (*η* = 0.001). Note that in order to train our model effciently we use batch update and one step of backpropagation to train 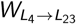 and we do not allow gradients to flow backwards in time.

Similarly, the learning rule for 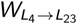 is given by the derivative of the cost function with respect to this weight matrix,

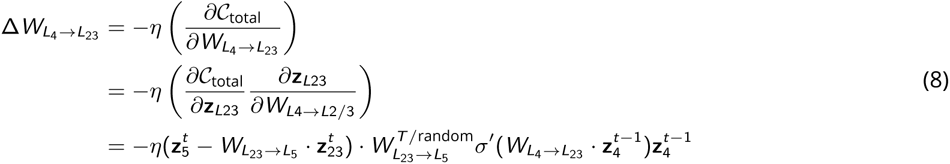

where 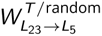 is the transpose of 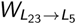 or a random matrix, depending on the experiments we performed (see Fig. 6).

Finally, the learning rules for *W*_Thal._*_→L_*_4_ and *W*_Thal._*_→L_*_5_ are given by,

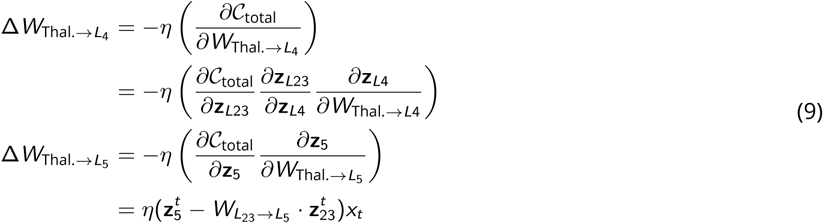

### Tasks

#### Gabor contextual-temporal task

In this task we generate synthetic sequential data. Each data point is a 28×28 Gabor patch with frequencies sampled from 𝒩(0.2, 0.1) with variability along x and y axis from 𝒰(3, 8) and fix orientations for each class with *θ* = [0, 18*^◦^*, 36*^◦^*,…, 162*^◦^*]. The top-down inputs can take values of [*−*18*^◦^*, 0*^◦^*, +18*^◦^*]. At each timestep, we randomly sample a datapoint *x_t_* with orientations *θ_i_* where *i* denotes the index number from the *θ* list, a top-down contextual input 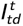 and generate the next input *x_t_*_+1_ by sampling a datapoint with the orientation 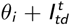. This setup means that for each input at timestep *t*, three subsequent orientations are possible, except angles 0*^◦^* and +162*^◦^*, which only have two possible successors.

This task is designed to investigate how Gabor patches at *t* can be predicted when their orientation is determined by a top-down variable. This top-down variable takes values of *−*18*^◦^*, 0*^◦^*, or +18*^◦^*, dictating how a Gabor patch in the recent past (*x_t−_*_1_) leads to a second (transformed) Gabor patch, *x_t_*). This results in sequences where the orientation shifts to the left, shifts to the right, or remains the same (see examples in Fig. 2b).

According to our model, at each timestep, the L5 network receives *x_t_* as its sensory input. Simultaneously, the L2/3 network processes the output of L4 (from previous timestep, *t −* 1) combined with the top-down contextual input.

#### Noise and occlusion tests

These experiments assess the model’s robustness to input degradation (Fig. 4). We focus on two forms of degradation: noise and occlusion. For noise, Gaussian noise is added to the input as 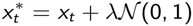, where *λ* scales the noise level (values from 0 to 1, in increments of 0.2). To examine the importance of the *L*2*/*3 *→ L*5 connection, we selectively disable the self-supervised cost during training. This prevents updates to L2/3 and L4 parameters through this loss, isolating the effects of this connection. Reconstruction performance across layers is measured using mean squared error between the reconstructed and the original (denoised) input.

For occlusion, random image sections are obscured with a dark patch (pixel values set to zero). After training, a Support Vector Machine is used to classify outputs of L2/3 and L5 based on the *x_t_* label. Classification accuracy on a held-out test set indicates how well the model copes with occlusions.

#### Visuomotor task

##### Simulating the Experimental Setup

To closely replicate the visuomotor task from Jordan and Keller ^19^, we have developed a method to generate synthetic sensorimotor data with modelled visual flow and motor speed. In our model, each vector dimension encapsulates a distinct aspect of visual flow, which is essential for simulating the sensory inputs typical in motion perception tasks. In particular, visual flow was calculated as **x***_t_* at any given time *t* following

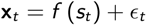

where, *f* denotes a function that converts speed into visual flow. We set *f* (*s_t_*) = *s_t_* to model a linear relationship (Fig. 7,8) or *f* (*s_t_*) = sin(*s_t_*) to model a non-linear interaction (Fig. S7c). The term *s_t_* represents the speed at time *t*. To mimic more realistic conditions we also add Gaussian noise, ϵ*_t_* = *𝒩*(0, 1). In our simulations, we model speed following a random walk. At each timestep, the speed *s_t_* is determined with equal probability between the following options

- Decreasing by 1: *s_t_* = *s_t−_*_1_ *−* 1
- Remaining the same: *s_t_* = *s_t−_*_1_
- Increasing by 1: *s_t_* = *s_t−_*_1_ + 1

This approach simulates the natural fluctuations of running speed, where an individual might slightly accelerate, decelerate, or maintain pace from moment to moment. After sampling speeds, we generate visual flow input *x_t−_*_1_ = *f* (*s_t−_*_1_) and *x_t_* = *f* (*s_t_*), with noise as defined above.

The speed variable provides top-down context, an important factor in our model that provides contextual information which help the model (L2/3) to predict the incoming visual flow given past visual flow and the current speed.

##### Mismatch simulation and measurement

After training, we obtain a baseline error signal by averaging the *prediction error* for each neuron across the dataset. Next, we simulate a mismatch (i.e. breaking the coupling between locomotion and visual feedback) by randomly setting the visual flow input to zero. Each mismatch period lasted for *K* timesteps (K=600 timesteps). We subsequently record the average prediction error signals for each neuron under simulated sensorimotor mismatches. To isolate the mismatch response for each neuron *z_i_*, we use the formula: *PE_i_* = *PE_i,baseline_* − *PE_i,mismatch_*. Here, *PE_i_* denotes the prediction errors, modeled as described below.

##### L2/3 and L5 prediction errors

The prediction errors for L2/3 and L5 that we analyse in Figs. 7 and 8 were calculate using the gradients of the cost function with respect to L2/3 and L5 neurons, respectively. Therefore, L2/3 prediction errors are calculated as:

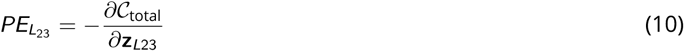

Whereas the L5 prediction errors are calculated as:

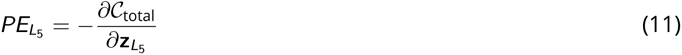

Note that because these prediction errors were calculated after learning converged, this means that the reconstruction cost was effectively zero.

### Sparsity metric

To measure activity sparsity of each layer in our model, we used the Treves-Rolls metric ^42,84^. The population sparseness, *S*, of each layer for a single stimulus was measured as:

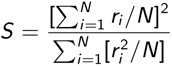

where *N* is the number of neurons, and *r_i_* the firing rate of neuron *i*. To get the average population sparseness for the entire sequence, we simply average *S* over a trial.

### Feedback and feedforward connection probabilities

In our work, we have tested the importance of feedback from L5 to L2/3 for learning. As part of this, we tested a range of connection probabilities *P* for this feedback pathway. The feedback connection from L5 to L2/3 are dropped with the probability of (1 *− P*_connectivity_, such that when the connection probability are set to 0, all the feedback connections are set to 0. The connection probability of all forward connections was set to 1.

## Data availability

Data to reproduce the simulated data can be generated using our code (see below).

## Code availability

The source code for the model proposed here and the respective analysis is available at https://github.com/neuralml/ neoSSL.

## Acknowledgements

We would like to thank the Neural & Machine Learning group, Randy Bruno, Nicol Harper, Andrew King and Katherine Willard for useful feedback. K.K.N. was funded by the UKRI Centre for Doctoral Training in Interactive Artificial, L.H. by the DFG (460088091), P.G.A by the Academy of Medical Sciences (SBF006\1047), Brain and Behaviour Research Foundation (YI Award 29300) and BBSRC (BB/X016331/1) and R.P.C. by the Medical Research Council (MR/X006107/1), BBSRC (BB/X013340/1) and a ERC-UKRI Frontier Research Guarantee Grant (EP/Y027841/1). We would like to thank Shuzo Sakata and Kenneth Harris for sharing their layer-specific sparsity data. This work made use of the HPC system Blue Pebble at the University of Bristol, UK. We would like to thank Dr Stewart for a donation that supported the purchase of GPU nodes embedded in the Blue Pebble HPC system.

## Author contributions

K.K.N. developed the computational framework with guidance from L.H. and R.P.C. K.K.N. performed all simulations and data analysis. K.K.N., P.A, L.H. and R.P.C. jointly wrote the manuscript. R.P.C supervised the project with help from L.H.

## Supplementary material

### Experimental details

We model each layer of the neocortical microcircuit as a linear transformation of their input followed by a non-linearity. The network layers are specified with the following default neuron counts: Layer 4 (L4) and Layer 2/3 (L2/3) each have 128 neurons, and Layer 5 (L5) has 16 neurons. In some experiments, we varied the number of neurons in each layer to explore the effect of neuronal density on the outcomes being evaluated. For experiments aligning with those conducted by Jordan and Keller ^19^, we adjusted the neuron counts in L2/3 and L5 to 32 and 16, respectively, to more closely match the proportion of neurons recorded from each layer during their experiments.

Network parameters are optimized using the standard ADAM optimizer with a learning rate of 0.001, and beta parameters (beta1 = 0.9, beta2 = 0.999). To achieve effcient learning, we employ standard stochastic gradient descent with a batch size of 32 and continue training until convergence, typically around 1000 epochs.

ANN parameters are initialized using an uniform distribution 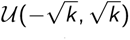, where *k* represents the number of neurons per layer. We use backpropagation for network training, except in experiments shown in Figure 6, where we explore alternative ways of setting the feedback weights. All results reflect an average across 5 random seeds.

Experiments were performed on the BluePebble supercomputer (University of Bristol), primarily using GeForce RTX 2080 Ti GPUs, with occasional CPU usage.

### L23-to-L5 feedback experiments

This experiment investigates the importance of the feedback connection from L5 to L2/3 in model performance. Instead of using the optimal feedback weight matrix 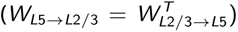, as derived using the backpropagation algorithm, we replace it with random weights in line with a variant of backpropagation known as feedback alignment ^48^. In addition, we introduce a probability variable (*P*) to control the density of the feedback connections (*W_L_*_5_*_→L_*_2_*_/_*_3_) compared to the forward connections (*W_L_*_2_*_/_*_3_*_→L_*_5_). Each feedback connection has a probability 1*− P* of being set to zero. For the results shown in Figs. 6 and S6 we created multiple model variants, each with varying feedback connection densities determined by this probability. All model variants are then trained on the sequential Gabor task.

## Supplementary figures

**Supplementary Figure S1.**
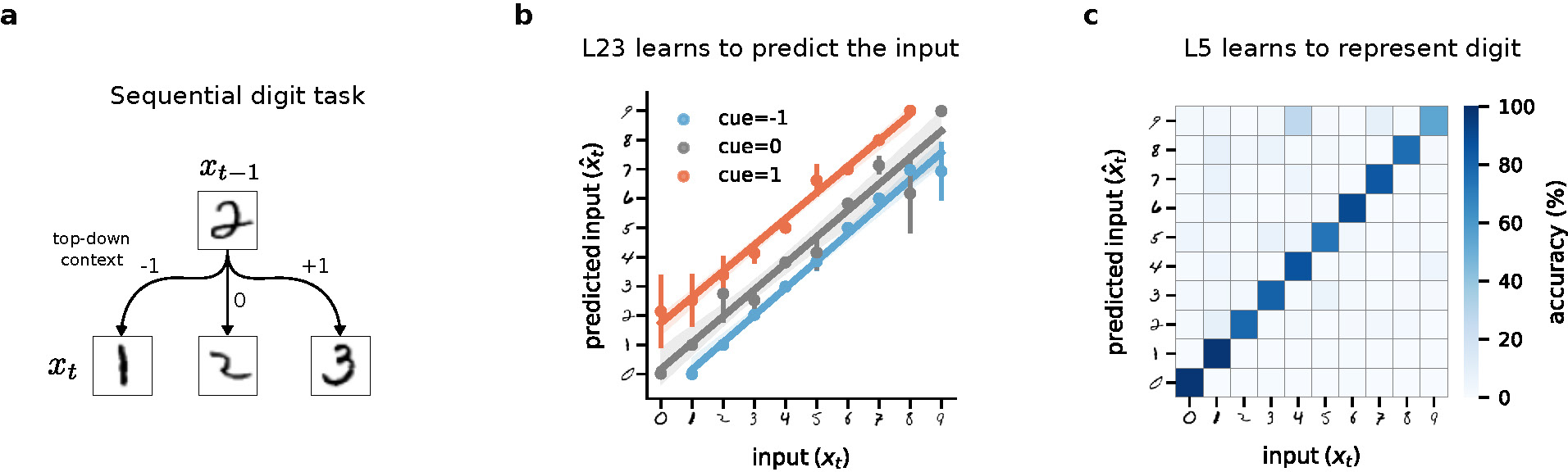
Learning a sequential digit task in a self-supervised canonical microcircuit. **a,** Sequential digit task used for training. The generative factor which is passed as top-down context determines the next digit. **b,** Prediction accuracy of a linear model trained on the output of L2/3. For a given input, L2/3 predicts the next possible input with high accuracy for all three cues (denoted by different colors). **c,** Confusion matrix showing the classification accuracy of a linear model trained on the output of L5.

**Supplementary Figure S2.**
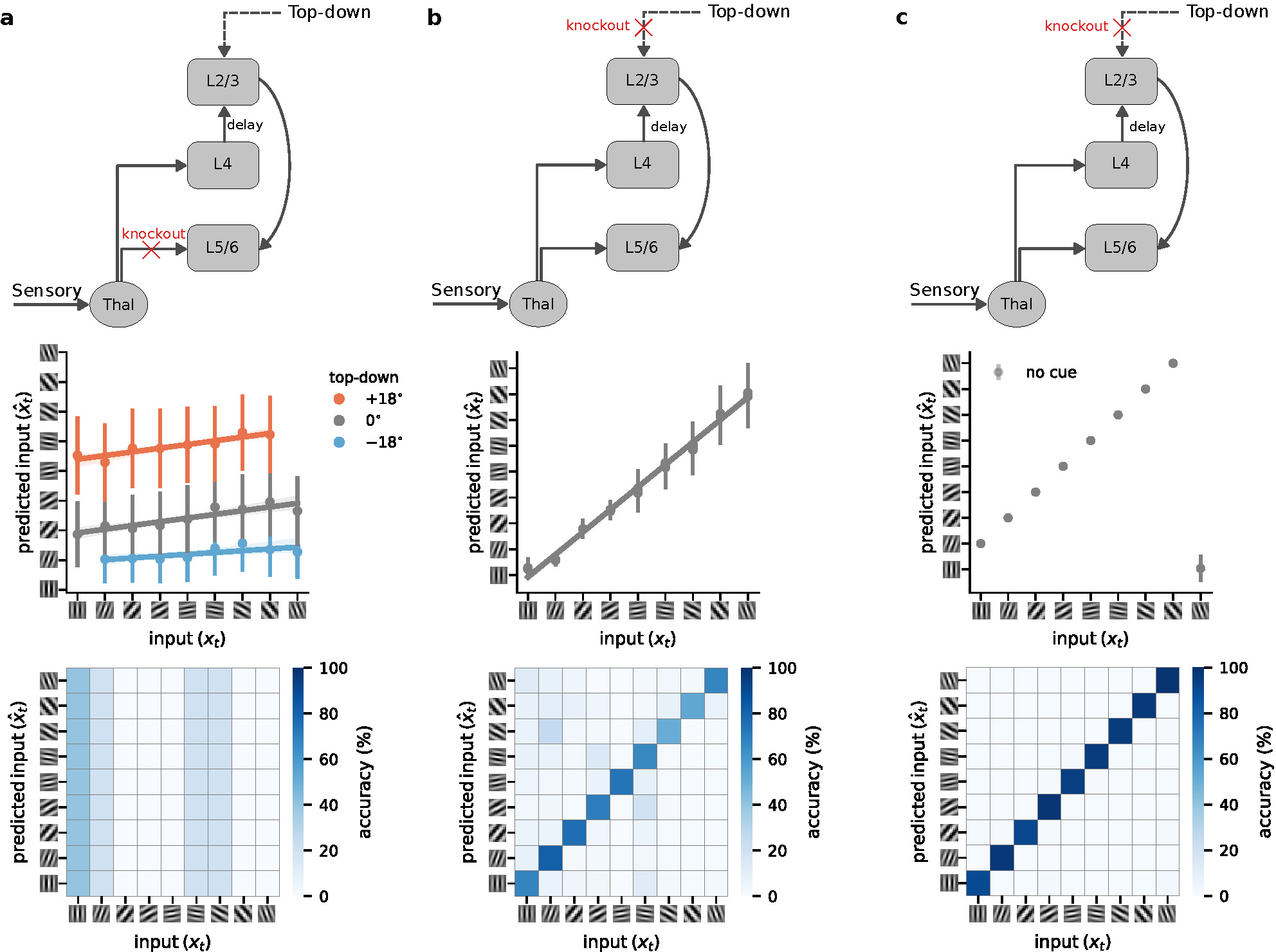
Prediction accuracy in L2/3 and L5 when connections are knocked-out. **a,** Connections from Thalamus to L5 are necessary for learning in both L2/3 and L5. **b,** Top-down input to L2/3 is crucial for L2/3 to predict the incoming input, while L5 performance is not significantly impacted when the top-down signal is deleted. **c,** Removal of top-down input to L2/3 in a deterministic task in which inputs always rotate clockwise (i.e. +18*^◦^*), top-down input is not required for task. In this case removing the top-down does not have any effect on the task performance.

**Supplementary Figure S3.**
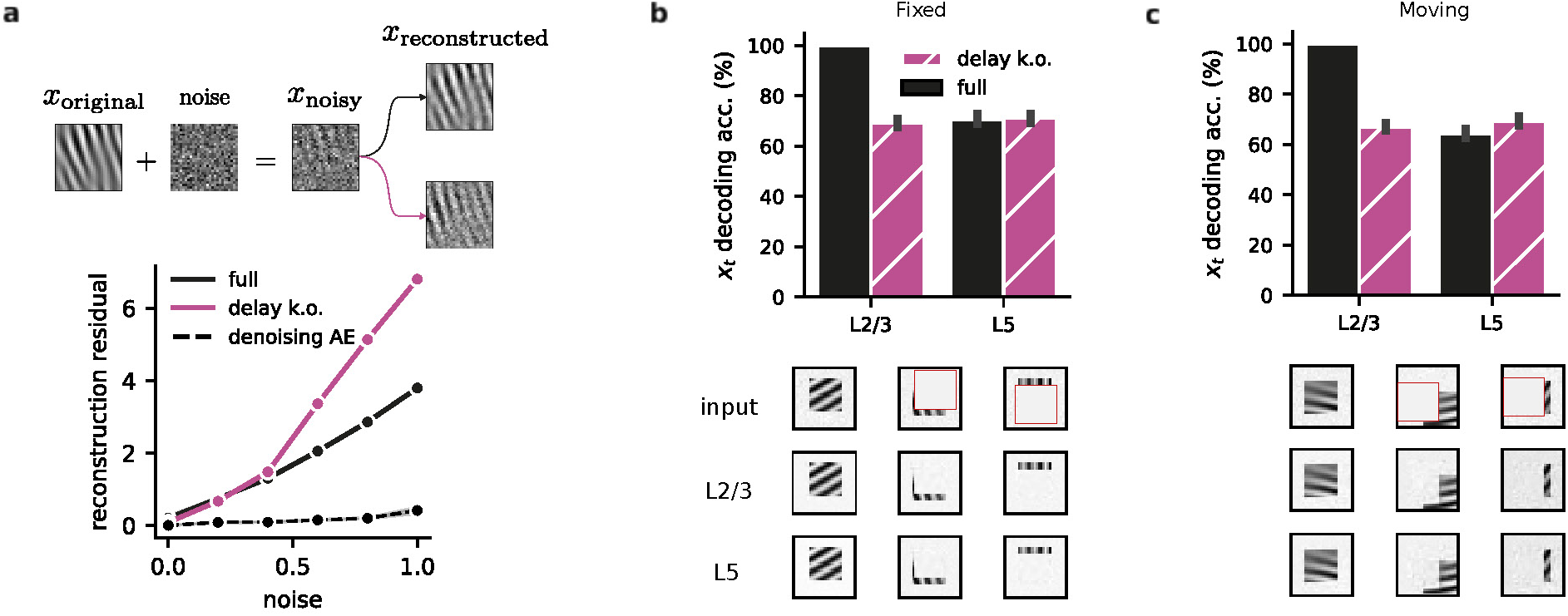
L4 to L2/3 delay increases denoising and robustness to occluded stimuli. **a,** L4 to L2/3 delay promotes noise suppression in L5 representations. Top: Schematic of noise added to the original inputs. Bottom: Noise-corrupted input samples lead to higher L5 reconstruction residuals (*x̂_t_ − x_t_*) when the L4-to-L2/3 delay is ablated (purple) compared to the full model (solid black). The dashed line represents the reconstruction residual for an autoencoder explicitly trained to denoise the input. **b,** Top: Decoding accuracy with and without L4 to L2/3 delay for a Gabor task with occlusion. Bottom: Three examples depicting L2/3’s losing it’s ability to recover occluded information, similar to L5’s incomplete reconstructions (top row: original occluded input; middle row: L2/3 reconstruction; bottom row: L5 reconstruction). **c,** Top: Accuracy with and without L4-to-L2/3 delay for a task in which Gabor patches move (top). Bottom: Examples further illustrate the robustness with moving Gabor patches (top row: original input with motion; middle row: L2/3 reconstruction; bottom row: L5 reconstruction).

**Supplementary Figure S4.**
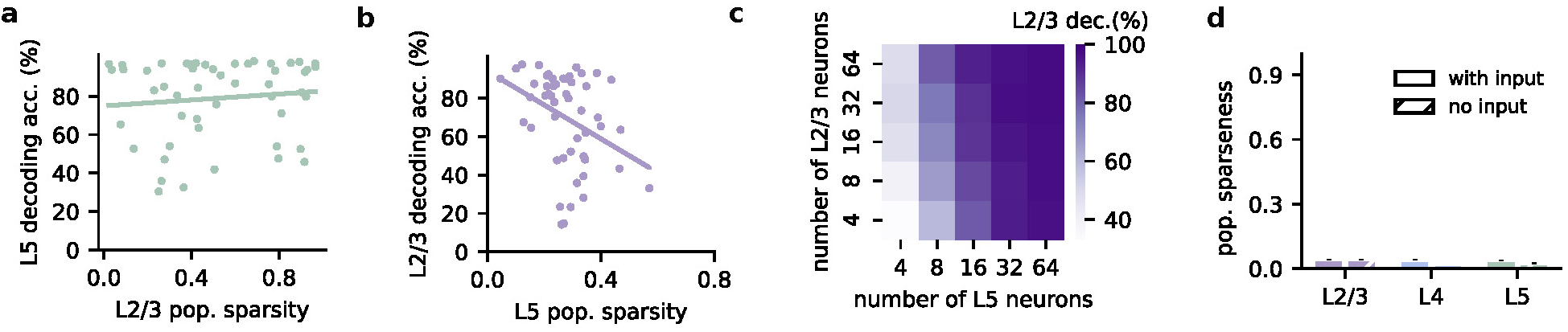
Sparsity and decodability of current input in layer 2/3 and layer 5. **a,** L5 decoding accuracy as a function of L2/3 population sparsity. **b,** L2/3 decoding accuracy as a function of L5 population sparsity. **c,** L5 decoding accuracy as a function of different numbers of neurons in L2/3 and L5. **d,** Effect of input removal on the sparsity of neocortical layers before training.

**Supplementary Figure S5.**
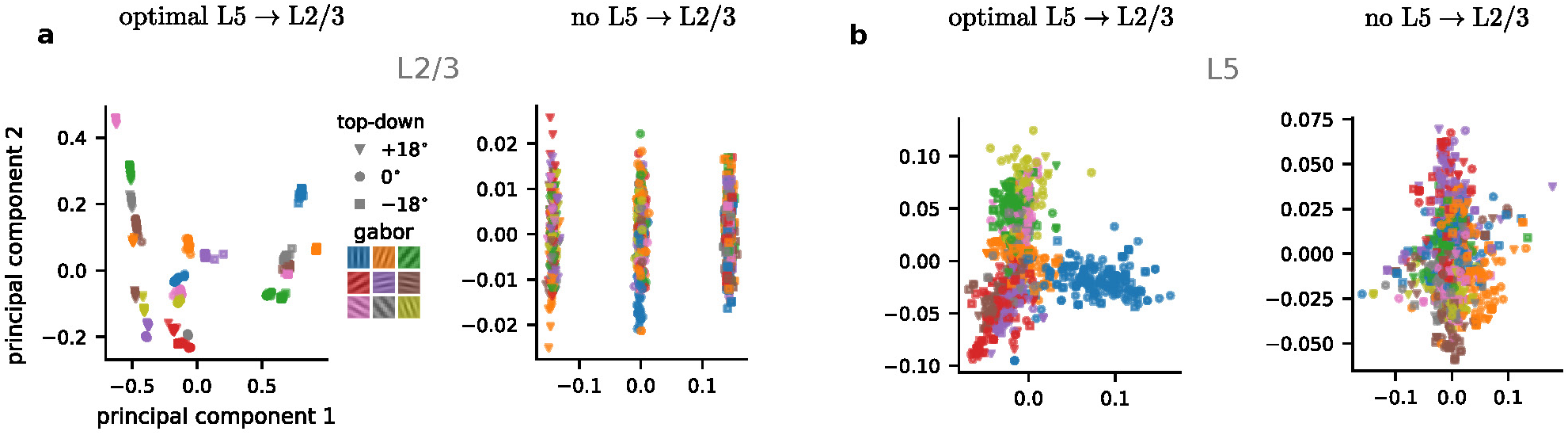
Low dimensional representations with optimal L5-to-L2/3 feedback in the Gabor temporal task. **a,** L2/3 representations are separated by context and Gabor orientation for optimal feedback (left) but are only grouped by context when the feedback is absent entirely (right). **b,** L5 representations between optimal feedback (left) and no feedback (right).

**Supplementary Figure S6.**
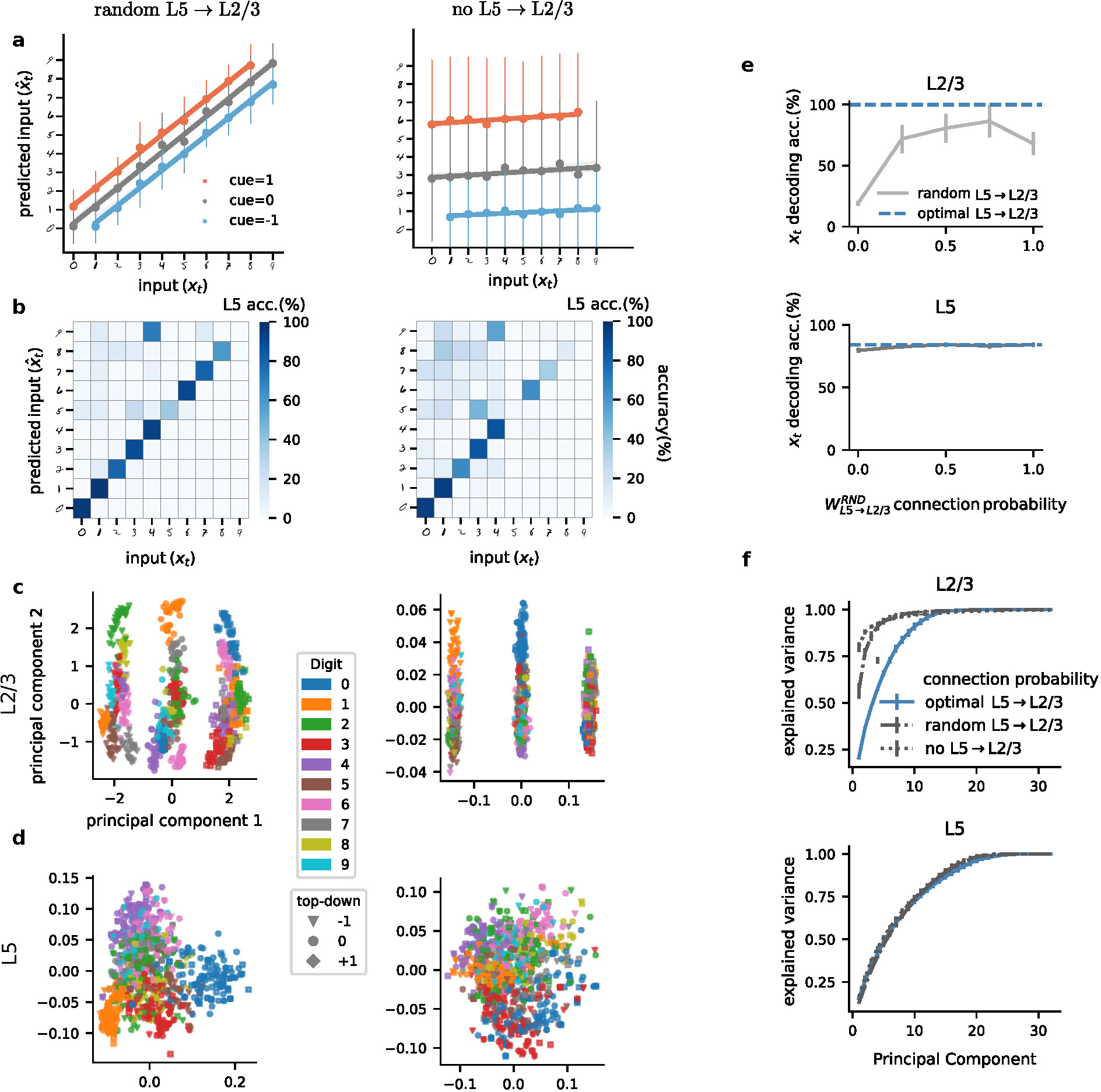
Role of feedback connections from L5 to L2/3 in learning a sequential digit task. **a,** L2/3 learns to predict the input with random feedback (left) but failed without feedback (right). **b,** L5 learns a good representation of the task with random feedback (left) and without feedback (right). **c,** L2/3 representations are separated by context and digit class for random feedback (left) but are only grouped by context when the feedback is absent entirely (right). **d,** L5 representation does not show a significant difference between random feedback (left) and no feedback (right). **e,** L2/3 linear prediction accuracy drops to chance level when the feedback is removed (top) while L5 classification accuracy is not impacted (bottom). **f,** The representation in L2/3 is mostly explained by the first few PCs for the ‘no feedback’ condition (top) while more PCs are required to explain the variance for the case with random feedback. L5 PCs explained variance is very similar in both conditions (bottom).

**Supplementary Figure S7.**
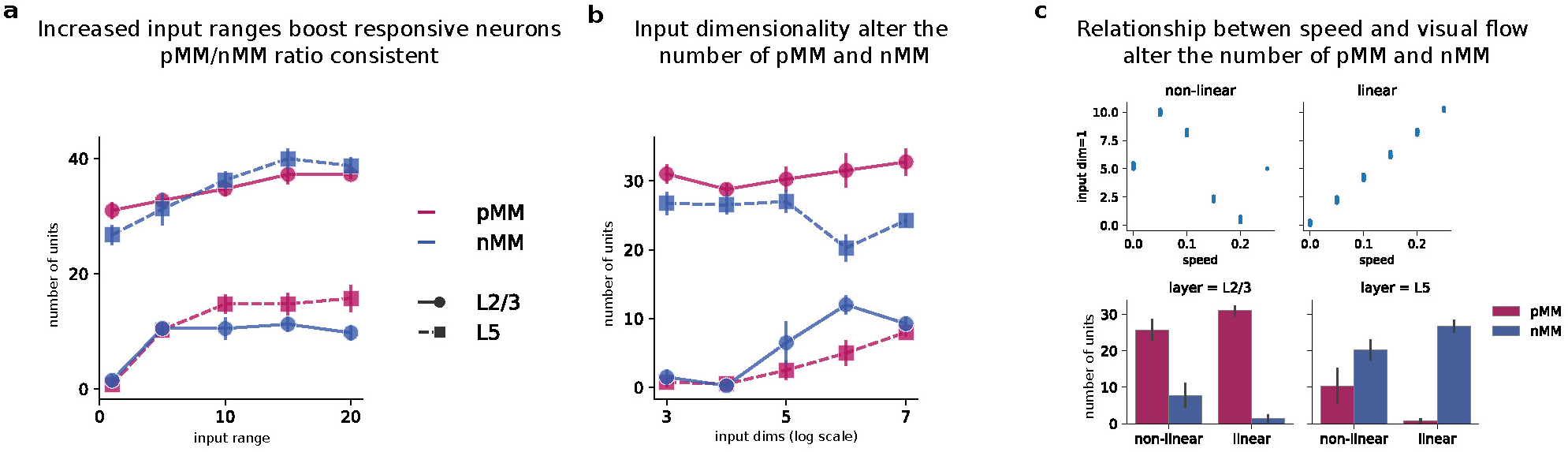
Mismatch errors across different model parameters. **a,** L2/3 and L5 prediction errors remain consistent across varying input values (pMM: positive mismatch errors; nMM: negative mismatch errors). **b,** L2/3 and L5 prediction errors are robust to the dimension of the input. **c,** L2/3 and L5 prediction errors persist with both linear and non-linear visual flow relationships to speed.

**Supplementary Figure S8.**
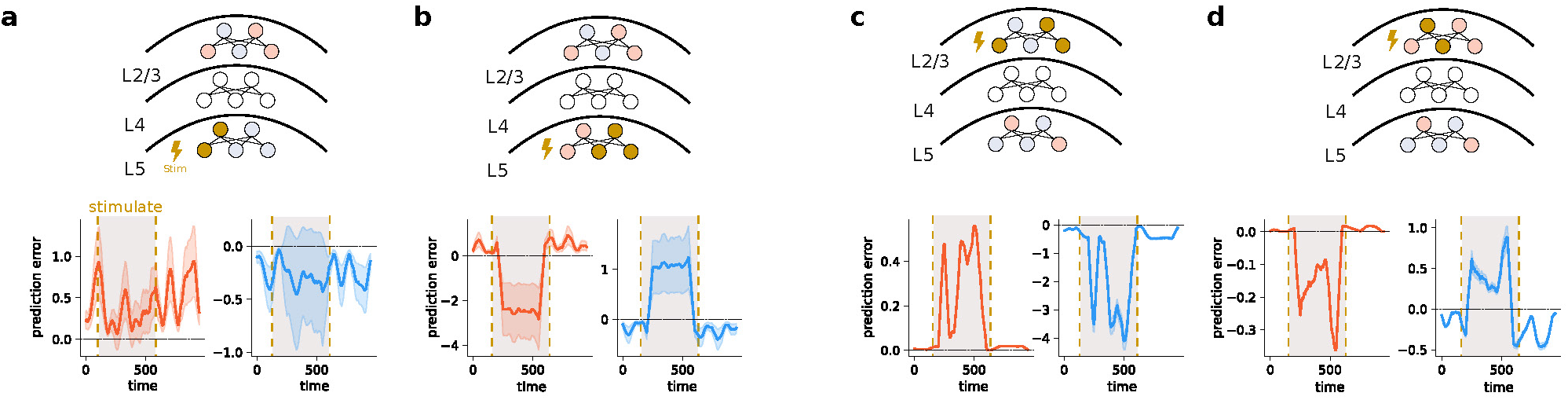
Mismatch errors in L2/3 and L5 after stimulating groups of neurons in L5 and L2/3, respectively. **a,** Increasing the activity of L5 neurons with positive errors during sensorimotor mismatch has variable effect on mismatch errors in L2/3. **b,** Modulating the L5 neurons with negative errors inverts the errors in L2/3. **c,** Scaling the output of L2/3 neurons with positive errors signals enhances the mismatch errors in both L5 neurons with negative and positive errors. **d,** Modulation of L2/3 neurons with negative mismatch errors flips the sign of errors in L5.

## Notes

https://github.com/neuralml/neoSSL

### Competing Interest Statement

The authors have declared no competing interest.

### Summary of Updates

Updated title and very minor changes in the paper.

